# Predation strategies of the bacterium *Bdellovibrio bacteriovorus* result in overexploitation and bottlenecks

**DOI:** 10.1101/621490

**Authors:** J. Kimberley Summers, Jan-Ulrich Kreft

**Affiliations:** Institute of Microbiology and Infection & Centre for Computational Biology & School of Biosciences, University of Birmingham, Edgbaston, Birmingham, B15 2TT, United Kingdom

**Keywords:** predator-prey interactions, generalist versus specialist, prey size, ordinary differential equations, mathematical modelling, robust permanence, competition, fitness

## Abstract

With increasing antimicrobial resistance, alternatives for treating infections or removing resistant bacteria are urgently needed, such as the bacterial predator *Bdellovibrio bacteriovorus* or bacteriophage. Therefore, we need to better understand microbial predator-prey dynamics. We developed mass-action mathematical models of predation for chemostats, which capture the low substrate concentration and slow growth typical for intended application areas of the predators such as wastewater treatment, aquaculture or the gut. Our model predicted that predator survival required a minimal prey size, explaining why *Bdellovibrio* is much smaller than its prey. A too good predator (attack rate too high, mortality too low) overexploited its prey leading to extinction (tragedy of the commons). Surprisingly, a predator taking longer to produce more offspring outcompeted a predator producing fewer offspring more rapidly (rate versus yield trade-off). Predation was only efficient in a narrow region around optimal parameters. Moreover, extreme oscillations under a wide range of conditions led to severe bottlenecks. A bacteriophage outcompeted *Bdellovibrio* due to its higher burst size and faster life cycle. Together, results suggest that *Bdellovibrio* would struggle to survive on a single prey, explaining why it must be a generalist predator and suggesting it is better suited than phage to environments with multiple prey.

**Importance:** The discovery of antibiotics led to a dramatic drop in deaths due to infectious disease. Increasing levels of antimicrobial resistance, however, threaten to reverse this progress. There is thus a need for alternatives, such as therapies based on phage and predatory bacteria that kill bacteria regardless of whether they are pathogens or resistant to antibiotics. To best exploit them, we need to better understand what determines their effectiveness. By using a mathematical model to study bacterial predation in realistic slow growth conditions, we found that the generalist predator *Bdellovibrio* is most effective within a narrow range of conditions for each prey. For example, a minimum prey size is required, and the predator should not be too good as this would result in over-exploitation risking extinction. Together these findings give insights into the ecology of microbial predation and help explain why *Bdellovibrio* needs to be a generalist predator.

## Introduction

Predator-prey relationships are some of the oldest and most important interactions in nature and occur at every level, from the smallest virus infecting a bacterium, to lions attacking wildebeest. Predators are often keystone species in natural ecosystems (1). Investigations into predator-prey dynamics in natural settings are, however, complicated by an array of confounding factors (2). Microbes by contrast make attractive models for studying predator-prey interactions in a controlled environment, with millions of individuals in a drop of liquid that can go through many generations in days. A completely separate, but increasingly important, reason for an interest in microbial predators is the antimicrobial resistance crisis, with resistance to even last resort antibiotics such as colistin rising (3). There is an urgent need for ‘living antibiotics’ as alternatives to antibiotics, such as phage and bacterial predators.

The best-studied bacterial predator is *Bdellovibrio bacteriovorus*. It is a Gram-negative bacterium that predates a wide range of other Gram-negative bacteria (4), regardless of their antimicrobial resistance or pathogenicity. *Bdellovibrio* alternates between two phases in its lifecycle. In the free-living attack phase, it does not grow but hunts prey. It swims faster than most bacteria, at speeds of up to 160 μm s^−1^ or ~100 body lengths s^−1^ (5). This requires a high metabolic rate and loss of viability within 10 hours if prey is not encountered (6). Once a cell is encountered, its suitability is investigated for several minutes (7). If suitable, it enters the prey’s periplasm by creating and squeezing through a pore in the outer membrane and peptidoglycan layer, shedding its flagellum in the process (8). Once inside the periplasm, *Bdellovibrio* kills the prey and lets the cytoplasmic nutrients leak into the periplasm (9). It also alters the prey’s peptidoglycan, causing the prey cell to round up into a “bdelloplast”. In this bdelloplast phase, *Bdellovibrio* uses these nutrients to grow into a long filament rather than dividing (5). When the nutrients have been used up, this filament septates into as many new cells as resources allow for, typically between 3 and 6 new predators per *E. coli* cell (10). If it were using normal binary fission, it could only produce 2^n^ offspring, potentially forsaking prey resources if they would only suffice for say 5 rather than 8 offspring. The new *Bdellovibrio* form flagella and lyse the remains of the prey cell, allowing them to burst out in search of fresh prey.

Mathematical models explain how predator-prey interactions can generate stable oscillations (11). Most models of microbial predator-prey interactions focussed on protists or bacteriophage. Only a few models considered predatory bacteria, see Table S1 for an overview of models and the review by Wilkinson (12). Varon & Zeigler (13) fitted a Lotka-Volterra model to their experimental results, which led them to propose that a minimal prey density is necessary for the survival of the predator. Crowley’s (14) model tracked substrate levels and used Monod kinetics for prey growth, it also included a delay between prey attack and the production of new predators to reflect the ~3 h long bdelloplast phase. The dynamics were destabilised by higher nutrient concentrations, especially at low dilution rates, leading to extreme oscillations. Wilkinson (15) developed two *Bdellovibrio* models, one based on a Holling type II functional response, but without specifically modelling the bdelloplast stage. The other model did include a combined predator-prey complex, but used a Holling type I functional response (15). Both models again showed a destabilising effect of nutrient enrichment, albeit somewhat ameliorated by the presence of a decoy species. Similarly, Hobley et al. (7), Baker et al. (16) and Said et al. (17) included a predator-prey complex and a Holling type I functional response. Hobley, Summers et al. (18) fitted a series of models, with a separate bdelloplast stage and various predator functional responses, to data from batch culture predation by either *Bdellovibrio*, a bacteriophage or both, and found a synergistic effect from dual predation. Two further models have been developed by Hol et al. (19) and Dattner et al. (20), both including spatial structure. These previous models focussed on the effects of dilution rate and substrate inflow on the dynamics of the chemostat systems. However, the effects of prey and predator characteristics such as prey size, attack kinetics and predation efficiency have not been studied.

In this study, we developed a family of models, based on ‘ingredients’ from previous models, to identify a unique combination of ingredients that generates realistic outcomes from realistic model assumptions and to ask many novel questions. This model predicts that *Bdellovibrio* has to be much smaller than its prey to survive, which is in agreement with empirical evidence (21). We also found that high attack rates, large prey sizes and low predator mortality, which might be expected to help the predator, are in fact detrimental for predator survival. Instead, these parameters have optimal values that maximize predator density. These optima occurred at the tipping points into oscillations and were often very narrow relative to the natural variation of these parameters. Moreover, we found that the system was prone to extreme oscillations in bacterial densities, leading to bottlenecks that would result in stochastic extinction. Additionally, *Bdellovibrio* would easily be outcompeted by a bacteriophage. Together, these three key predictions (narrow optima, population bottlenecks and phage superiority) suggest that *Bdellovibrio* would struggle to survive on a single prey species in a natural environment full of phages, and where conditions are unlikely to be optimal, and if so, not for long. Our findings are consistent with the fact that *Bdellovibrio* is a generalist predator, so we posit that our model explains why *Bdellovibrio* is a generalist predator and why it has to be as small as it is.

## Model development

We developed models to investigate the effects of a consumer, the predatory bacterium *Bdellovibrio bacteriovorus* or a bacteriophage, on a bacterial population under continuous culture conditions in a chemostat. A chemostat captures the low substrate concentrations and slow growth rates typical of the natural environment (22). Also, the prey on its own will reach a steady state population density in the chemostat, which facilitates the investigation of oscillations caused by predation. Oscillations could not be observed in a batch culture model. We developed a family of models to investigate the effect of changes in the structure of the model on the predator-prey dynamics. This structural sensitivity analysis (23) identified Model 6 as the most realistic in terms of assumptions and predictions (see SI text, Table S2 and Fig. S4), so we explored Model 6 further and describe it in Fig. 1 and list its parameters and their values in Table 1. Model 6 included a single abiotic resource (substrate S), single prey species (N) and a single obligate predator (P). The prey species grew by consuming the substrate according to Monod kinetics. The predator had a Holling type II functional response (predation rate proportional to predator density but saturating at high predator density), a bdelloplast (for *Bdellovibrio*) or infected cell (for bacteriophage) stage and mortality. The bdelloplast (B) is a distinct stage in the *Bdellovibrio* life cycle that usually lasts for 2 to 4 hours. There are distinct parallels between this bdelloplast stage and a bacteriophage infected cell as the prey cell does not grow and replicate when infected by a bacteriophage or consumed by *Bdellovibrio* and the predator or phage does not prey on or infect further prey. We modelled the bdelloplast or infected cell stage as a separate entity to account for the delay in producing offspring and the combining of the consumer and its prey. For simplicity, these entities will be referred to as a bdelloplast from now on, but the principles apply equally to an infected cell.

**FIG 1.**
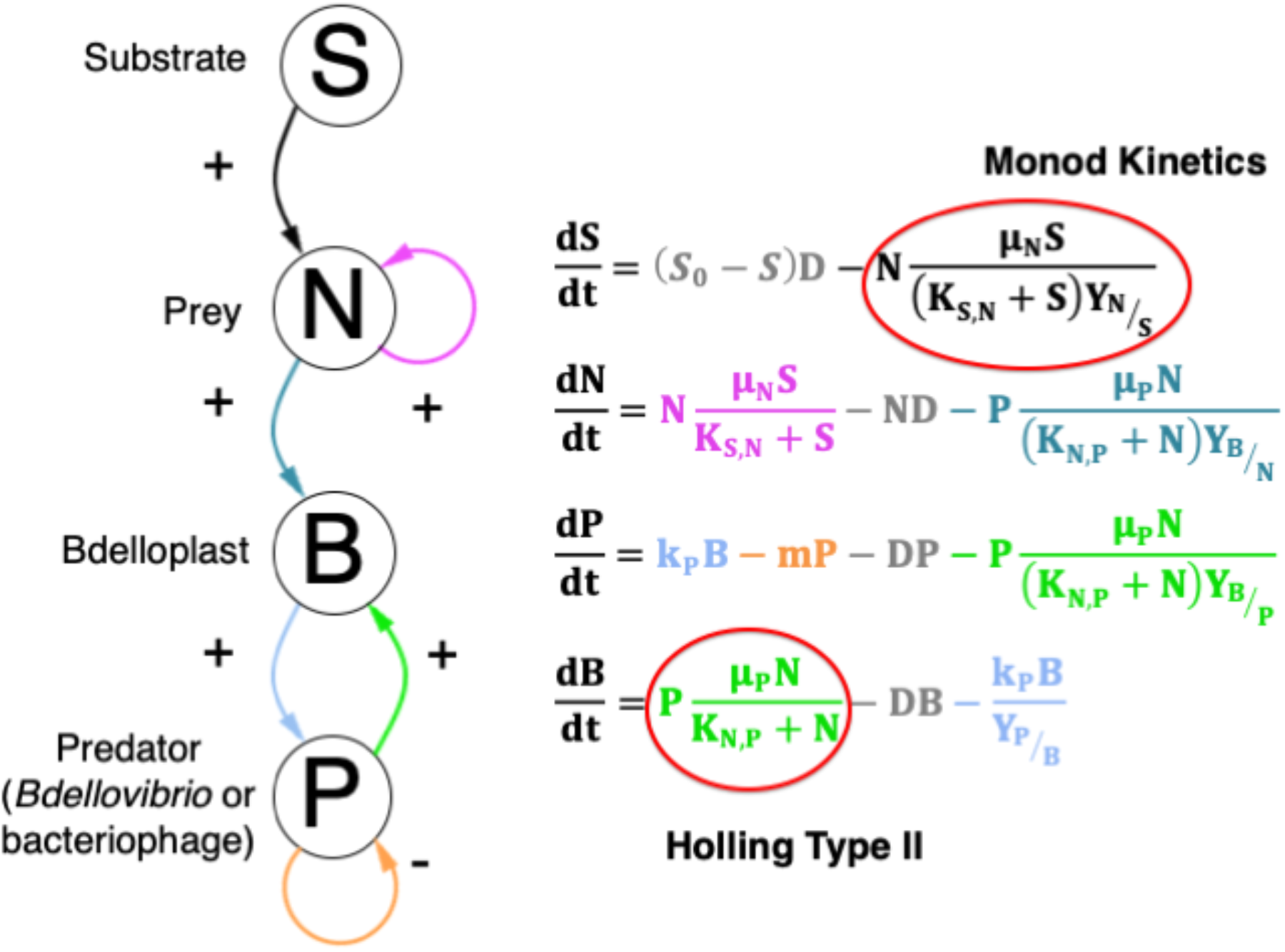
Principal model used to track predator and prey densities under chemostat conditions (Model 6, see Table S2). Substrate is consumed by prey to fuel growth. Prey and predators combine to form a bdelloplast. The bdelloplast matures to give new predators. Predators have mortality. This is the model used unless stated otherwise.

**Table 1.** Baseline parameters for the principal model (Model 6) and range over which global sensitivity analysis was performed. All yields are expressed in the form 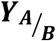, which is the yield of consumer A per resource B consumed. The half-saturation constant, or *K*-value, of consumer B for resource A is expressed as ***K***_***A,B***_.

| Parameter | Units | Value | Minimum | Maximum | Source |
| --- | --- | --- | --- | --- | --- |
| Inflow substrate concentration ($S_0$) | mg ml<sup>-1</sup> | $5 \times 10^{-3}$ | $5 \times 10^{-3}$ | 2.5 | Assumed to be $\sim 2 \times K_{S,N}$ |
| Prey maximum specific growth rate ($\mu_N$) | h<sup>-1</sup> | 1.23 | $\frac{3}{4} \ln(2)$ | $3 \ln(2)$ | (24) |
| Predator attack rate constant ($\mu_P$) | mg bdelloplast<br>mg predator<sup>-1</sup><br>h<sup>-1</sup> | 0.38 | 0.3 | 6 | (18) |
| Predator $K$-value ($K_{N,P}$) | mg prey<br>ml<sup>-1</sup> | $8.6 \times 10^{-4}$ | $1.5 \times 10^{-6}$ | $6.5 \times 10^{-3}$ | (18) |
| Yield of bdelloplasts from predators ($Y_{B/P}$) | mg bdelloplast<br>mg predator<sup>-1</sup> | 8 | 5 | 20 | Relative sizes of *E. coli* and *Bdellovibrio* cells (4, 25) |
| Bdelloplast maturation rate ($k_P$) | mg predator<br>mg bdelloplast<sup>-1</sup><br>h<sup>-1</sup> | 0.109 | 0.075 | 0.3 | (10) |
| Predator mortality rate ($m$) | h<sup>-1</sup> | 0.06 | 0.03 | 0.09 | (6) |
| Dilution rate ($D$) | h<sup>-1</sup> | Always 0.05 | | | |
| Prey $K$-value ($K_{S,N}$) | mg<br>ml<sup>-1</sup> | Always $2.34 \times 10^{-3}$ | | | (24) |
| Yield of prey from substrate ($Y_{N/S}$) | mg dry mass<br>prey<br>mg substrate<sup>-1</sup> | Always 0.4444 | | | (24) |
| Yield of bdelloplasts from prey ($Y_{B/N}$) | mg bdelloplast<sup>-1</sup><br>mg prey<sup>-1</sup> | Always $\frac{Y_{B/P}}{Y_{B/P} - 1}$ | | | Relative sizes of *E. coli* and *Bdellovibrio* cells (4, 25) |
| Yield of predators from bdelloplasts ($Y_{P/B}$) | mg predator<br>mg bdelloplast<sup>-1</sup> | Always 0.438 | | | Relative sizes of *E. coli* and *Bdellovibrio* cells and burst size (4, 10, 25) |

## Results

### Model implementation and validation showed the system to be brittle

We first ensured that the numerical results were reliable by testing all MatLab Ordinary Differential Equation (ODE) solvers. Only the Runge-Kutta ode45 solver could correctly handle all test cases including extreme oscillations (Fig. S1). It was also necessary to set tolerances of this solver to very low values and to constrain variables to be non-negative (Fig. S2). Secondly, we used the same test cases to compare simulations using biomass-based units with those using particle-based units, required to obtain numerically stable simulations with bacteriophage parameters. The results were in agreement (SI text and Fig. S3). These unusual difficulties of obtaining correct numerical results highlight that the system tends to undergo extremely rapid changes, followed by periods of stasis, and is very sensitive to parameter settings.

### Structural sensitivity analysis identified most appropriate model

To gain a better understanding of the impact of various modelling choices, we compared a number of ordinary differential equation (ODE) and delay differential equation (DDE) models (Table S2, Fig. S4). In all cases, the same set of standard conditions, based on *Bdellovibrio* predating *E. coli* that are growing on glucose (0.05 mg ml^−1^) and a dilution rate of 0.0333 h^−1^ (equivalent to a 30 hour retention time) were used. We first ran the simulation without predators until the prey had reached a steady state (same for all models). (Model 1) Addition of a predator without a bdelloplast stage (or other form of delay between prey killing and predator birth) gave rise to sustained, extreme oscillations. (Model 2) Incorporating an explicit delay of 4 hours, approximately doubled the oscillatory period from ~150 hours to ~300 hours. (Model 3) Adding mortality reduced the oscillatory period to approximately 100 hours. (Models 4-6) Addition of an explicit bdelloplast stage stabilised the system, resulting in a stable steady state of co-existing predator and prey, regardless of the predator’s Holling type functional response. (Model 4 vs. 6) With a Holling type I predator functional response (Model 4), the final prey density was lower, and the predator density higher, than with the saturating type II response (Model 6). (Model 5) Constant input of prey to the system; this is akin to growing the prey on its own in one chemostat that feeds into a second chemostat containing predator. This gave a very similar response to the single chemostat with constant input of substrate. Model 6 was chosen as the most biologically appropriate model for all further work, for several reasons. Firstly, the time required for *Bdellovibrio* to consume the contents of a prey cell, convert these into new *Bdellovibrio* predators, septate and finally lyse the prey cell to release new predators is about four hours (10). As the prey is killed very shortly after penetration (26), there is a significant delay between prey killing and birth of new predators, which is best modelled by treating the bdelloplast stage as a separate entity. Secondly, *Bdellovibrio* has a high endogenous respiration rate and a correspondingly low life span in the absence of suitable prey (6). Hence, it is appropriate to include predator mortality in the model. Thirdly, the saturating Holling type II functional response results from the ‘handling time’ of a predator (27). This would correspond to the minimum time for a successful attack in a prey-saturated environment, i.e., the attachment and penetration time of *Bdellovibrio* of ~10 minutes (28).

### Dynamic regimes from steady states to extreme oscillations

Model 6 has 12 parameters and six possible dynamic regimes (Fig. 2a). In order to gain a better understanding of the factors determining which regimes were observed, we swept through a range of inflow substrate concentrations (*S*_0_) and dilution rates and evaluated the steady state and its stability analytically (see model analysis in SI). As expected, at the highest dilution rates and lowest *S*_0_, all biological species washed out. Reducing the dilution rate or increasing *S*_0_ enabled survival of first the prey alone and then the predator. Further increases in *S*_0_ destabilised the system, resulting first in damped and then sustained oscillations, before finally reaching a linearly unstable state. We ran simulations in each of the regimes and found that they agreed with the outcomes predicted from the model analysis (Fig. 2b-g), apart from giving sustained, extreme oscillations where the analytical results predicted a linearly unstable state that would correspond to washout (Fig. 2d).

**FIG 2.**
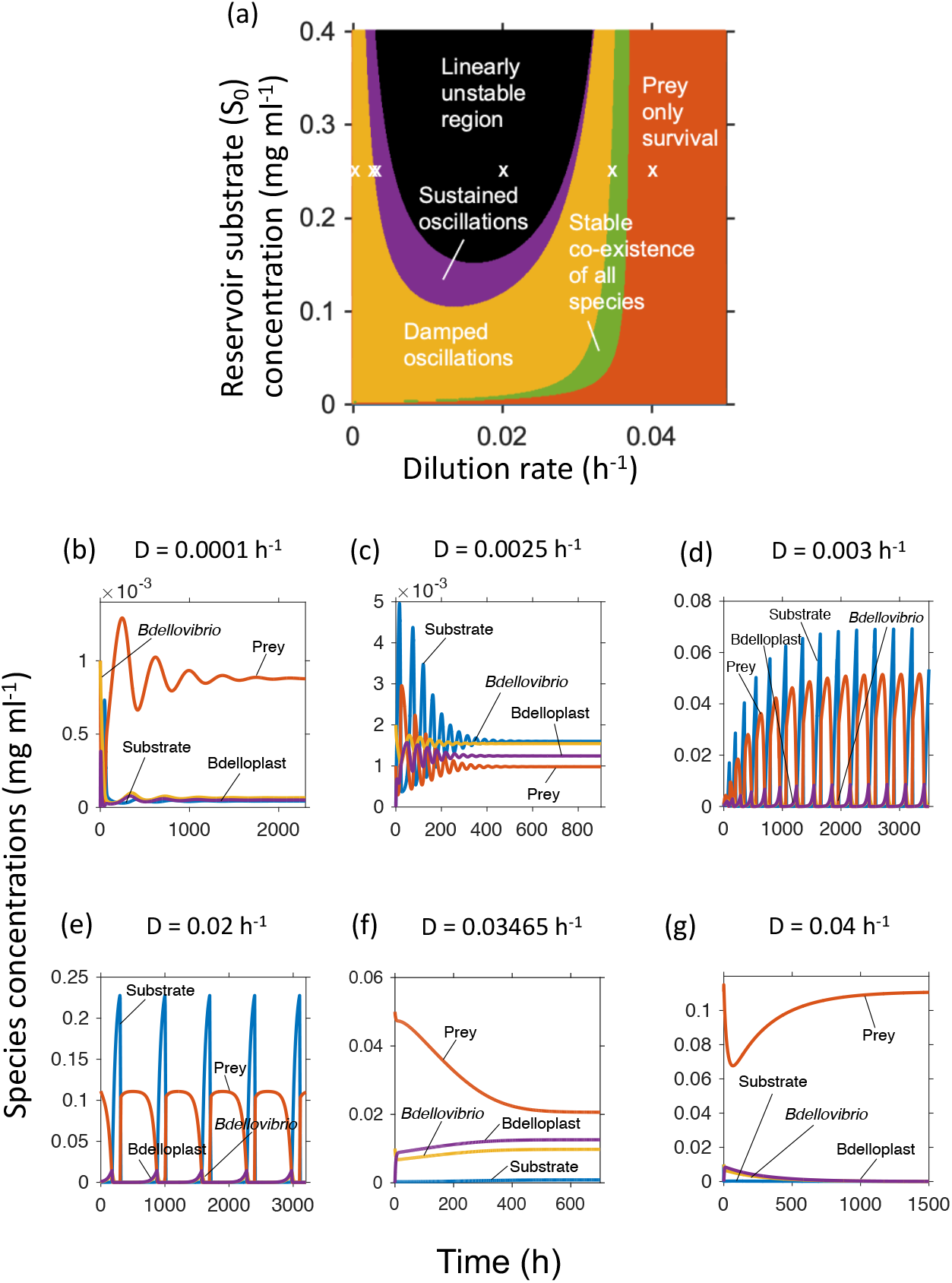
Dynamic regimes of Model 6. **a** Analytically calculated regimes, as depending on the inflow substrate concentration *S*_0_ and dilution rate *D*. **b**-**g** Simulations at *S*_0_ = 0.25 mg ml^−1^ and increasing dilution rates, indicated by white crosses in **a**. **b** Damped oscillations that ended in steady state co-existence. **c** Damped oscillations of much shorter period than **b**. **d** Amplifying oscillations that ended in sustained, extreme oscillations. **e** The analytically predicted linearly unstable region gave sustained, extreme oscillations in the numerical simulations. **f** Stable co-existence of predator and prey. **g** Predator extinction.

Since the oscillations generated by our *Bdellovibrio* model had a much longer period, extremely abrupt rises and falls, and a strong asymmetry in wave shapes compared to the typical Lotka-Volterra models for animal populations, we compared our model with a protist predator model by Curds and Bazin (29) that generated similarly extreme, but shorter period, oscillations. The period of the protist model could be increased to that of the *Bdellovibrio* model just by replacing protist with *Bdellovibrio* kinetic parameters (SI text on Protist model, Fig. S6).

Dimensional analysis was then performed to deduce which combinations of parameters determine the qualitative behaviour of the system, yielding 7 independent parameter combinations. Essentially, rate parameters became relative to the dilution rate; inflow substrate concentration relative to the prey’s *K*-value and the predator’s *K*-value replaced by its ratio to the prey *K*-value (for details see SI text). These *K*-values are half-saturation constants and a low *K*-value corresponds to high affinity and vice-versa. The next 7 sections show the effect of these 7 dimensionless parameter combinations that determine qualitative behaviour.

### Bdellovibrio has a minimal and optimal attack rate constant

To our knowledge, Varon & Zeigler (13) and Hobley, Summers et al. (18) are the only studies of predation kinetics where experimental data were used to infer parameters of a model. Varon & Zeigler (13) used a relative of *Bdellovibrio*, the marine strain BM4, and *Photobacterium leiognathi* as prey. *P. leiognathi* can grow at rates of up to 0.2 h^−1^ (30), however, nothing beyond the study by Varon and Zeigler (13) is known about the growth kinetics of strain BM4. Hobley, Summers et al. (18) used the type strain of *Bdellovibrio bacteriovorus* (HD100), with *E. coli* as the prey species, which are the species assumed in our study. Predictions from the Hobley, Summers et al. study were used to calculate default values for the attack kinetics. Since different predator and prey combinations may have different kinetics, we investigated the effect of varying the attack rate constant (*μ*_*P*_), which in the Holling type II functional response corresponds to the catalytic rate constant of an enzyme with Michaelis-Menten type saturation. The ratio of attack rate constant and *K*-value (*K*_*N,P*_) gives the initial slope of the Holling type II function, which determines the predation rate at low prey densities. We expected that the faster the predator was at locating and attacking prey, the more successful it would be, particularly given its high mortality (half-life of 10 hours), implying that *Bdellovibrio*’s attack rate constant was limited by intrinsic constraints preventing it from being any faster (at up to 160 μm s^−1^ it is already one of the fastest swimming bacteria known, especially considering its small size). Instead, we found that the highest attack rate constant was not the best, as a lower value was optimal for the abundance of the predator, and conversely for the prey (Fig. 3). The position and width of this optimum varied with dilution rate (Fig. 3c, d). Increasing levels of *S*_0_ narrowed the range of *μ*_*P*_ in which the predator could achieve near maximal density. Below the optimal *μ*_*P*_, there was a sharp drop to predator extinction. The optimal *μ*_*P*_ was also the rate at which the system underwent a Hopf bifurcation (31) from a stable steady state of co-existence into an oscillatory regime (Fig. 3e-g). The finding that too fast predation was not optimal can be understood by realising that the food source for the predator, a living organism, is a renewable resource, if it is given time to regrow. Hence, a too effective predator reduces regrowth of its prey and will starve.

**FIG 3.**
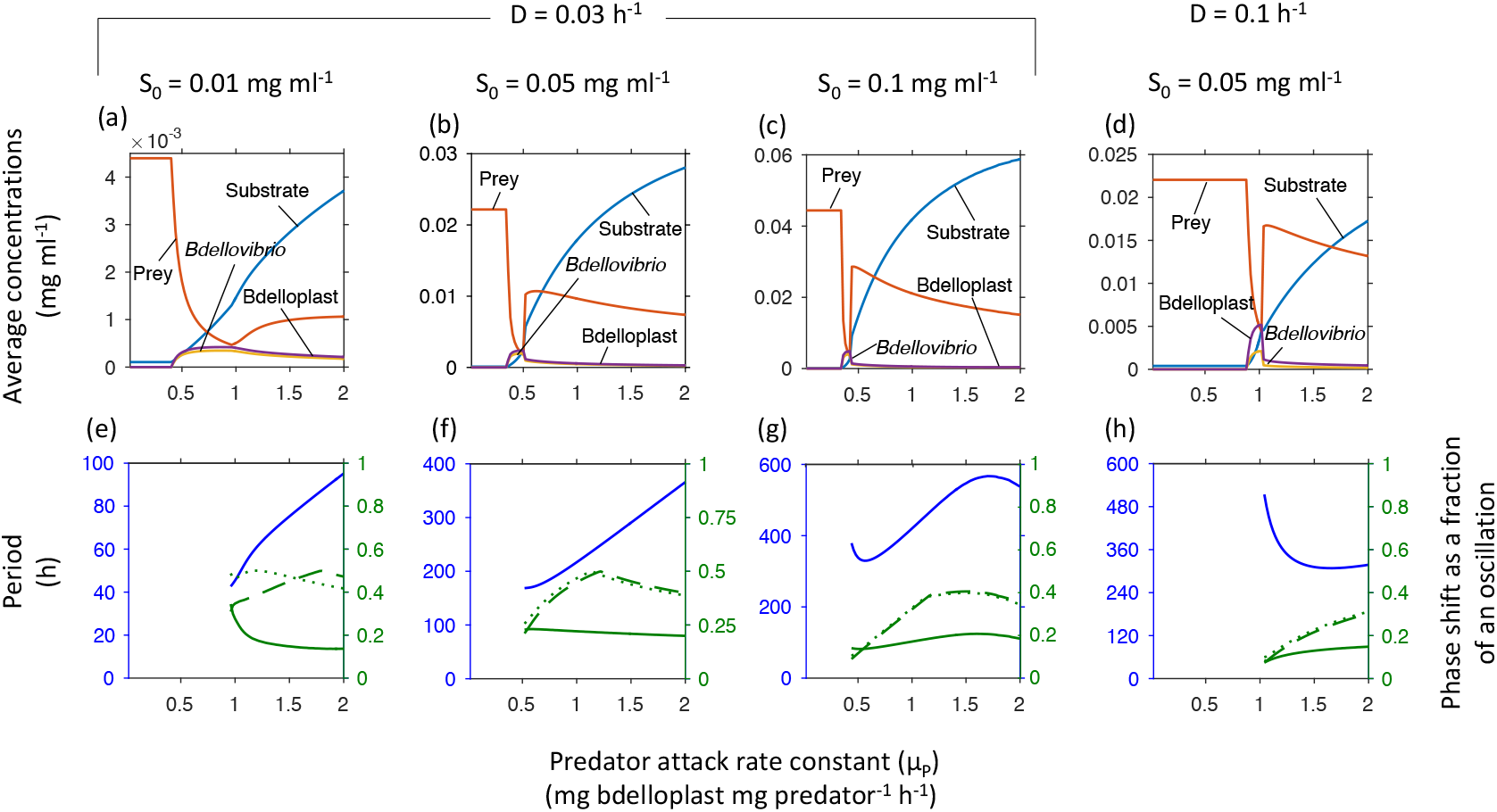
Minimal and optimal attack rate constant (*μ*_*P*_). The average population densities and substrate concentrations, oscillatory periods and phase shifts strongly depend on *μ*_*P*_, shown at increasing inflow substrate concentrations (*S*_0_) and two dilution rates (cf. phase diagram in Fig. 2a). Top row shows concentrations at steady state or averaged over one oscillatory cycle. Bottom row shows the oscillatory period (blue, left axis) and phase shifts (green, right axis) from substrate peak to peak of prey (solid line), free *Bdellovibrio* (dashed line) or bdelloplast (dotted line). Note that oscillations start above the optimal *μ*_*P*_. In order to obtain accurate simulation results at all parameter values, the absolute tolerance of the ode45 solver had to be reduced from 1 × 10^−9^ to 1 × 10^−12^.

### Higher prey growth rate does not benefit prey

Since *Bdellovibrio* can prey on a wide range of Gram-negative bacteria, which have a wide range of maximum specific growth rates (*μ*_*N*_), not just *E. coli*, we investigated the effects of prey growth rate. We expected that an increased prey growth rate would benefit both predator and prey. Instead, increasing prey growth rate benefitted only the predator and never the prey. At low inflow substrate concentration (*S*_0_), populations were co-existing in a stable steady state, where surprisingly prey growth rate had no effect on prey and predator density (Fig. 4b, cf. phase diagram in Fig. 2a). At high *S*_0_, populations were co-existing in sustained oscillations, and increasing prey growth rate led to decreasing prey abundance and then a sharp drop in prey abundance at the bifurcation point where the system became stable (Fig. 4a, c). Overall, higher prey growth rate did not benefit the prey.

**FIG 4.**
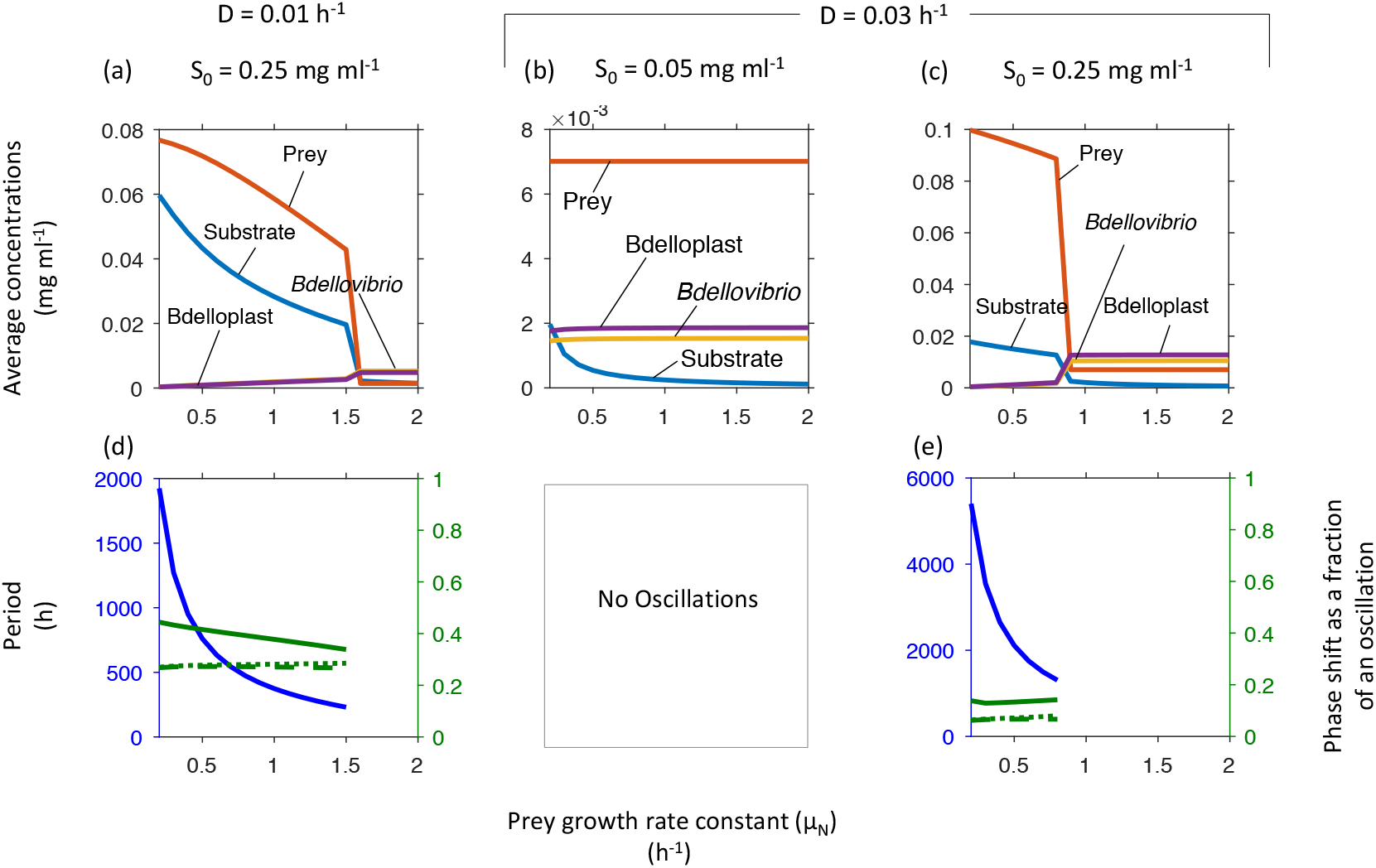
Increasing the maximal specific growth rate of the prey (*μ*_*N*_) leads to a sharp drop in prey density and stabilizes the system at high inflow substrate concentrations (*S*_0_). The system was more stable at low *S*_0_ (cf. phase diagram in Fig. 2a). Top row shows concentrations at steady state or averaged over one oscillatory cycle. Bottom row shows the oscillatory period (blue, left axis) and phase shifts (green, right axis) from substrate peak to peak of prey (solid line), free *Bdellovibrio* (dashed line) or bdelloplast (dotted line). Note that oscillations occur at the higher *S*_0_ and below a critical *μ*_*N*_. Prey density is much higher in the oscillatory regimes, where it decreases with increasing *μ*_*N*_. Since prey density will of course be 0 if *μ*_*N*_ is zero, there is an optimal *μ*_*N*_.

### Predators benefit from mortality

Mortality of *Bdellovibrio* is much higher than that of other bacteria. Therefore, we included it in our model and swept death rates from 0 to 0.2 h^−1^ – substantially more than the 0.06 h^−1^ reported for *Bdellovibrio* (6). Surprisingly, predator death was beneficial for the predator with an optimal death rate just above a critical mortality where oscillations were replaced with stable co-existence (Fig. S7). Further increases in mortality caused predator extinction. Increasing inflow substrate concentration (*S*_0_) and dilution rate (*D*) narrowed the predator peak, giving a narrow window for predator persistence (Fig. S7).

### There is an optimal maturation rate for bdelloplasts

The bdelloplast stage, where the predator grows inside the prey periplasm with a certain specific growth rate (maturation rate - *k*_*P*_) until prey resources are exhausted and offspring is produced, takes ~3 hours with *E. coli*. We found, as with most other parameters, that there was a minimal maturation rate required for predator survival, and an optimal maturation rate (Fig. S8). This optimal rate was just below a critical rate, above which populations oscillated at higher inflow substrate concentration.

### Highest affinity of the predator for its prey is not always optimal

The dimensional analysis identified that the system behaviour depends on the ratio of the *K*-values *K*_*N,P*_ and *K*_*S,N*_ (SI text, so we swept through a range of values). At low inflow substrate concentrations (*S*_0_), populations did not cycle and the lowest *K*_*N,P*_ was optimal (Fig. S9a). At high *S*_0_, in contrast, too low *K*_*N,P*_ resulted in oscillations with reduced average predator levels (Fig. S9b-e). Raising the *K*_*N,P*_ resulted in a bifurcation from extreme oscillations to a stable co-existence that benefited the predator at the expense of the prey. The optimal *K*_*N,P*_ for the predator was just above this critical *K*_*N,P*_ (SI text).

### Increasing prey productivity benefits predator until an optimum is reached

Substrate inflow (*S*_0_) determines prey productivity. Increasing prey productivity from 0 at the outset benefits the predator much more than the prey, which remains at very low levels, until a predator maximum is reached. Further increasing substrate inflow led to a drop in predator and rise in prey (SI text, Fig. S10).

### Bdelloplast burst size

The last of the 7 dimensionless parameter combinations is the burst size, which increases with prey size. It is not surprising that there was a minimal burst size for predator survival, but it was not expected that this minimal burst size (at higher dilution rates or inflow substrate concentrations) was higher than the experimentally determined burst size for *E. coli* of 3.5 (Fig. S11). We also found an optimal burst size, after which predator abundance declined, i.e., too large prey was not optimal. The lower the dilution rate, the lower the minimal and optimal burst size and the broader the optimum such that the optimum was in the range corresponding to typical prey sizes (Fig. S11). Increasing the burst size above the optimal value resulted in a bifurcation to extreme oscillations, corresponding to a sharp rise in prey and drop in predator average densities, but only at low *S*_0_. At higher *S*_0_, the optimum was very narrow, above values corresponding to typical prey and oscillations did not occur (Fig. S11).

### Bacteriophage outcompetes Bdellovibrio

We asked whether bacteriophages would outcompete *Bdellovibrio* on single prey populations, and if so, under which conditions and why. The bacteriophage was parameterised based on the well-studied T4 phage infecting *E. coli*. T4 caused oscillations and outcompeted *Bdellovibrio* (Fig. 5). Why did the phage win? There are three processes where the two predators differ. Firstly, the attack kinetics, where the phage had a higher attack rate constant (*μ*_*P*_), but a higher *K*-value (*K*_*N,P*_). Secondly, the kinetics of prey consumption, where the phage had a higher burst size 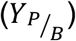 and a faster maturation rate (*k*_*P*_). Thirdly, the phage, unlike *Bdellovibrio*, did not have mortality. To find out which advantage(s) allowed the phage to win, we ran competitions where the phage kept the prey *K*_*N,P*_ disadvantage and had one or more of the advantages. Increased burst size 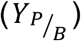 alone was sufficient for the phage to win (Fig. S12d). A combination of increased *μ*_*P*_ and reduced mortality was also sufficient (Fig. S12a, b, e). Increased *k*_*P*_ was insufficient even in the presence of either an increased *μ*_*P*_ or reduced mortality (Fig. S12c, f, g). Simulating the experiments of Williams et al. (32) where a bacterial predator (but no phage) responded upon addition of prey to a mesocosm, suggests that if suitable phage were present, they should have responded better to the prey addition (SI text, Fig. S13).

**FIG 5.**
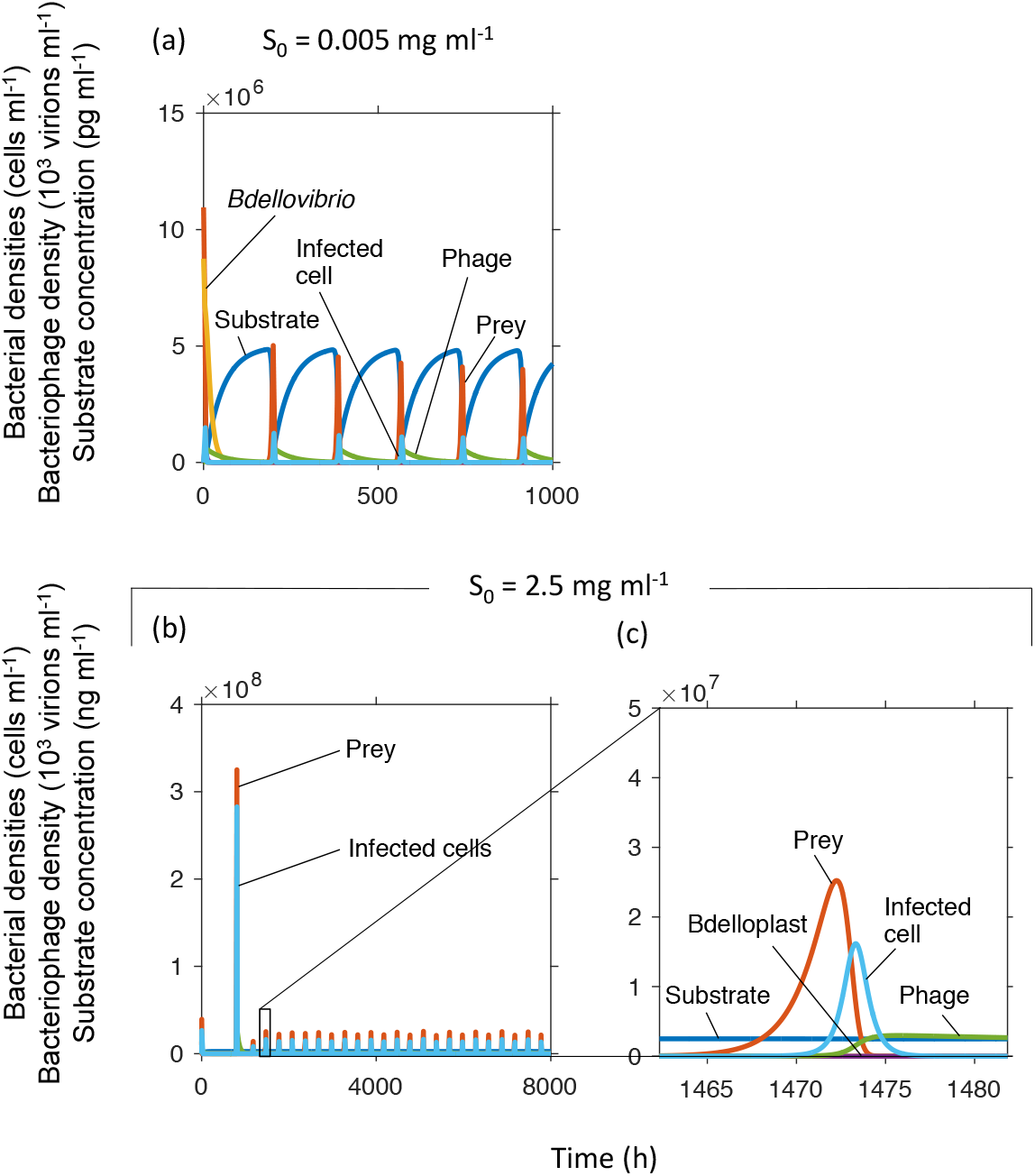
T4 phage caused oscillations and outcompeted *Bdellovibrio* at two very different inflow substrate concentrations. Dilution rate was 0.02 h^−1^. Note that units had to be changed to particle densities from the biomass densities used in the other sections. Panel c shows a zoomed in version of a peak in panel b, the substrate concentration appears to be constant on the scale needed to show the other variables.

### Trade-off between high rate versus high yield for predators

Since it takes time to convert prey resources into predator biomass, more complete exploitation of prey resources (higher yield or burst size) should come at the cost of a longer maturation time before offspring will emerge (lower maturation rate - *k*_*P*_). The fast but wasteful predator was assumed to convert bdelloplasts into offspring at a higher rate (*k*_*P*_ one third higher), but had a lower burst size (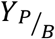 halved) to reflect a reduced yield. The high yield strategy outcompeted the high rate strategy (Fig. 6). Note that the phage had both higher burst size and faster maturation rate, considering this, it is not surprising that the phage won (cf. Fig. 5).

**FIG 6.**
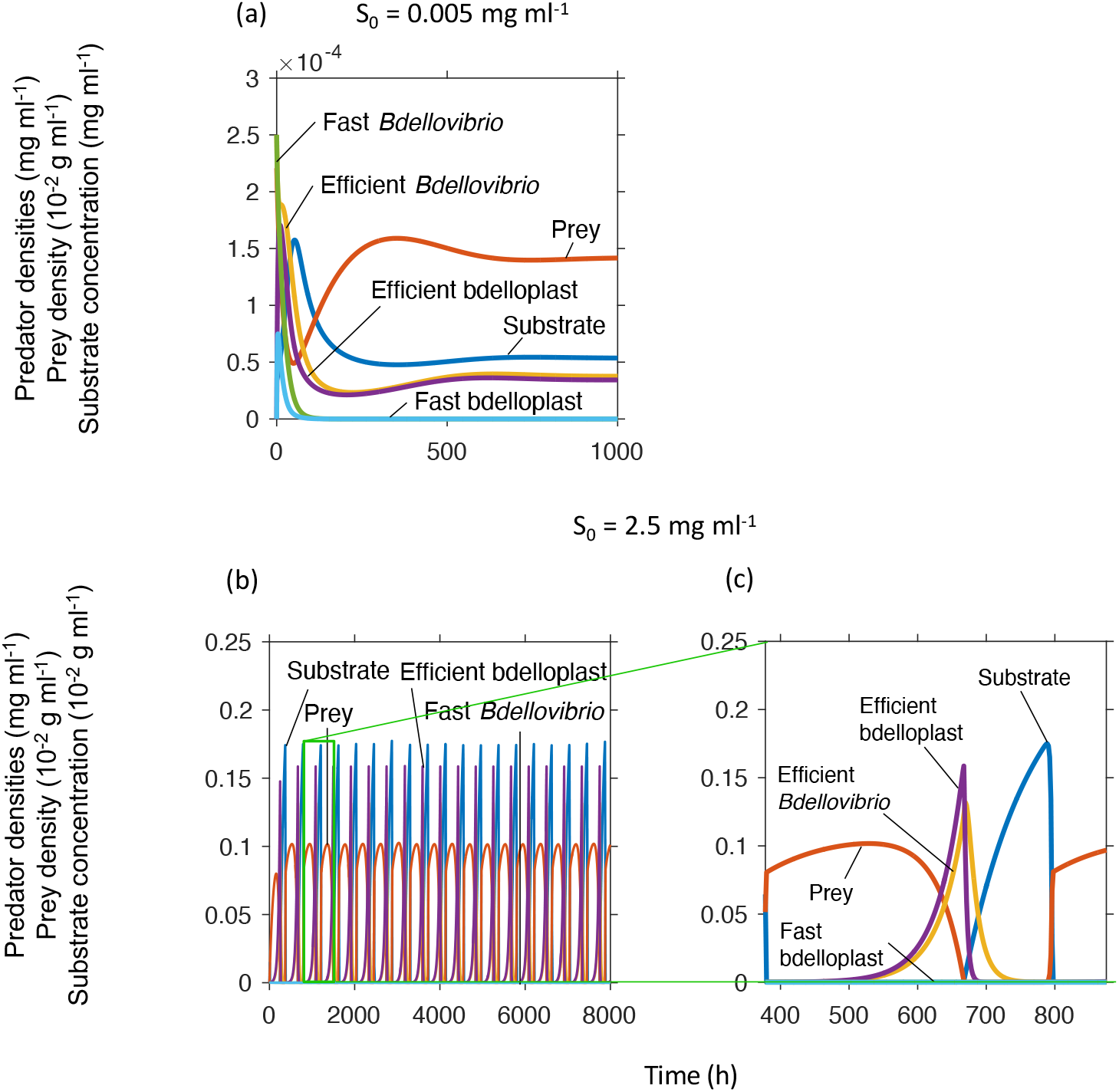
A slow, but economical (efficient resource use, therefore high 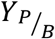) predator outcompetes a fast (high *k*_*P*_) but wasteful predator at different inflow substrate concentrations (*S*_0_). Dilution rate = 0.02 h^−1^, (a) *S*_0_ = 0.005 mg ml^−1^ and (b) *S*_0_ = 0.25 mg ml^−1^. Panel (c) shows a zoomed in version of a peak of panel (b).

### Co-existence of two predators on one prey

We assumed that a trade-off between attack rate constant (*μ*_*P*_) and *K*-value (*K*_*N,P*_) exists because time has to be invested for finding prey as well as for binding and examining potential prey. We found that this trade-off would allow coexistence of a ‘fast’ predator with a higher *μ*_*P*_ and a ‘high affinity’ predator with lower *K*_*N,P*_ on a single prey (Fig. S14).

### Minimal and optimal prey cell sizes

Bacteria have a large range of sizes (and therefore biomass) (33). Clearly prey can be physically too small to be entered and consumed. *Bdellovibrio* also needs to produce at least two offspring per prey cell. Cells of *B. bacteriovorus* are around seven times smaller than *E. coli* cells; this small size might enable *Bdellovibrio* to prey on cells with a wider range of sizes. While consuming a larger prey cell will produce more offspring, it will take longer. Also, larger prey cells will mean fewer prey are available if prey growth is limited by the substrate entering the system as in our chemostat case and most environments. It is not obvious whether the higher number of offspring per prey would offset the disadvantages of longer maturation times and fewer cells to hunt. We found that there was both a minimum prey to predator size ratio required for predator persistence and an optimal value for maximal predator biomass (Fig. 7). Fat prey caused the system to display extreme oscillations. The optimal prey size was just below the size causing these oscillations. Increases in *S*_0_ narrowed the optimum towards the minimal prey size (Fig. 7a-c). Increases in dilution rate also narrowed the peak, but increased the optimal prey size (Fig. 7c, d).

**FIG 7.**
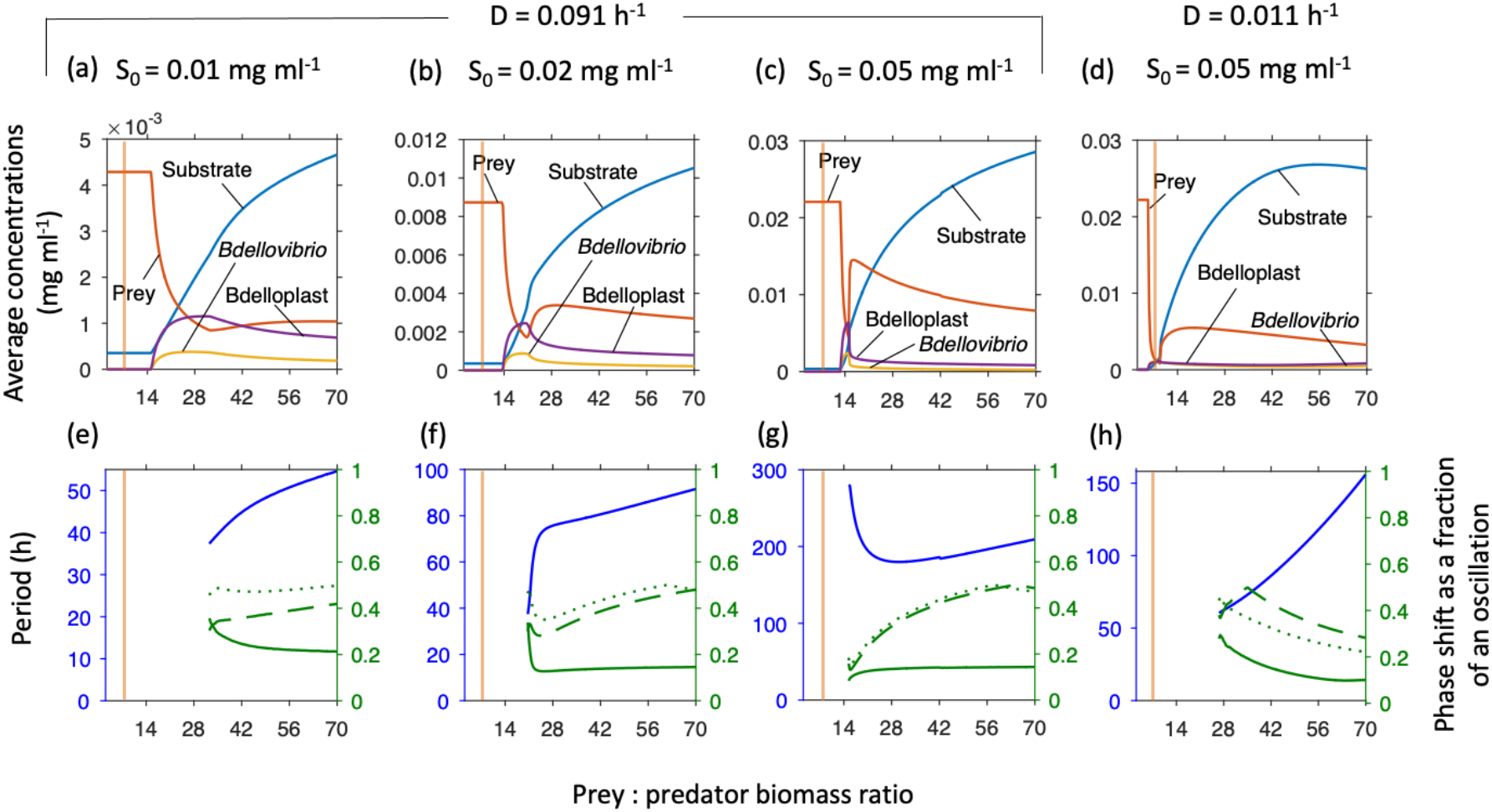
Minimal and optimal prey biomass, relative to predator biomass (0.028 pg dry biomass predator cell^−1^). Predator density showed a broad optimum at low inflow substrate concentrations (*S*_0_) that became narrower at higher *S*_0_ and dilution rate. Too large prey caused oscillations. Top row shows concentrations at steady state or averaged over one oscillatory cycle. Bottom row shows the oscillatory period (blue, left axis) and phase shifts (green, right axis) from substrate peak to peak of prey (solid line), free *Bdellovibrio* (dashed line) or bdelloplast (dotted line). The vertical orange line, at a prey:predator ratio of 7, is the biomass ratio between an average *E. coli* and *Bdellovibrio* and similar to most prey used in laboratory studies or for isolating *Bdellovibrio* from the environment.

### Bottlenecks, permanence and paradox of enrichment

When sweeping parameters, it became clear that the system oscillated with long periods of hundreds of hours and extreme amplitudes for many of the conditions tested, with extremely low minimum values (< 1 × 10^−30^ mg dry mass ml^−1^). Such bottlenecks would result in the extinction of the predator (or both predator and prey) in any natural system prone to extrinsic fluctuations and governed by stochastic processes. We therefore examined how prey size (previous section) in combination with dilution rate and inflow substrate concentration (cf. Fig. 2a) would affect robust predator persistence (union of regions of stable co-existence and damped oscillations in Fig. 2a). Robust persistence is known as permanence and means that population densities do not approach zero as the boundary is a repeller (34). The prey size range allowing permanence narrowed with increasing *S*_0_ (Fig. S15). This effect of increased productivity (higher influx of resource for prey) destabilising the system is known as the paradox of enrichment (35) and is typical for predator-prey systems. Prey also had to be unrealistically large to enable permanence at higher dilution rates and lower productivity.

### Tragedy of the commons

Similarly to increasing productivity, the prey size range enabling permanence shrank with increasing attack rate constant of the predator (Fig. S16). This is an example of the tragedy of the commons, where overexploitation of a shared resource known as the commons is to the detriment of the users of the resource, such as overfishing (36). Here, a too effective predator that consumed its prey faster than it could regrow led to extinction of the predator. Increases in the inflow substrate concentration (*S*_0_) narrowed the prey size range for permanence further whilst increases in dilution rate expanded the range (Fig. S16).

### Global parameter sensitivity analysis

To understand how much the system behaviour would change if the parameters were different due to changes in substrate, prey or predator species, we conducted a global sensitivity analysis (SI text and Figs. S17, S18). Substrate concentration was only sensitive to the inflow substrate concentration. Prey density was not sensitive to substrate concentration and prey growth kinetics, but sensitive to predator parameters. Predator densities were sensitive to the attack rate constant, but also to the maximal specific prey growth rate.

## Discussion

Our results suggest that *Bdellovibrio* consuming a single prey species can only survive permanently within a very narrow range of conditions. We therefore characterise this predator-prey system as ‘brittle’. For example, over a wide range of conditions, the system is prone to extreme oscillations with periods of over a hundred hours and bacterial densities dropping below 0.1 pg ml^−1^. In a deterministic mathematical model, species densities can eventually recover from even the smallest positive number but in a biological system, there has to be at least a single cell left (~1 pg). More importantly, when a system contains just a few cells over a long time, stochastic fluctuations will almost inevitably result in the loss of those few cells, leading to local extinction. For most system parameters, there was a narrow range of values that allowed the predator to reproduce fast enough to avoid being washed out and not trigger these oscillations, and the optimal value to maximise predator numbers occurred near the threshold triggering these oscillations. Increasing nutrient concentrations narrowed this survival range. Previous models of microbial predator-prey interactions in chemostats, including those with *Bdellovibrio* as the predator, also showed a tendency to extreme oscillations, especially for higher nutrient concentrations (14, 15, 29, 37). Only the group of Varon studied this experimentally in chemostats with the *Bdellovibrio* like predator BM4 and *Photobacterium leiognathi* as prey. They did observe oscillations, albeit less extreme ones than in our model (38, 39). They also found that BM4 could survive in a chemostat with a prey density of 2-5 × 10^4^ CFU ml^−1^ (39), substantially less than the 7 × 10^5^ CFU ml^−1^ predicted by their model (13) and less still than the 4.4 × 10^6^ CFU ml^−1^ minimum required for survival in our model. In contrast, Keya and Alexander (40) found *Bdellovibrio* strain PF13, isolated from soil, would only replicate in the presence of at least 3 × 10^7^ CFU ml^−1^ of its *Rhizobium* prey. Studies so far do not allow us to judge whether predictions are reasonable.

Given the brittle behaviour predicted by the model it is surprising that *Bdellovibrio* is ubiquitous in non-marine environments, while *Bdellovibrio* like organisms are ubiquitous in marine environments (41–44). This suggests that there must be other forces at work stabilising population numbers. One possibility is that *Bdellovibrio* is a hotspot organism targeting habitats of high prey density, or structured environments containing areas of high prey density such as biofilms, and that these hotspots are the sources of *Bdellovibrio*, and other habitats are sinks. Biofilms represent a lump of prey for *Bdellovibrio*. It possesses the lytic enzymes needed to chew the extracellular matrix that holds cells within the biofilm and can likely derive valuable nutrients from this (45). *Bdellovibrio* can predate the metabolically inactive cells found deeper within biofilms and its gliding motility allows it to move within the biofilm (46). *Bdellovibrio* is also highly motile, and contains chemotaxis genes that might enable it to locate these prey rich areas (47) – so hotspots that are only temporary can be located. Indeed, there is ample evidence that *Bdellovibrio* is a hotspot organism. It moves towards regions rich in bacteria (48). Higher numbers are found in sewage rather than in rivers (49, 50), and downstream sewage treatment plants rather than upstream (51) or in sewage treated rather than untreated soils (52). The prey rich rhizosphere sustains higher predator numbers than the bulk soil (53). Eutrophic lakes support higher numbers than oligotrophic lakes (54). *Bdellovibrio* like organisms are more abundant in marine sediments than the water column and much higher in oyster shell biofilms (55). *Bdellovibrio* are likewise more common in trickling filter biofilms than in their inflow (56).

However, the model predicts a narrower range of conditions for survival at the higher nutrient concentrations expected in hotspots. Therefore, it is unlikely that these hotspots would sustain *Bdellovibrio* if it were only predating a single prey species. Moreover, our model also suggests that bacteriophage would outcompete *Bdellovibrio* under all conditions tested. This is because phage, as specialist predators, have several big advantages. Indeed, bacteriophage are far more numerous in nature than *Bdellovibrio* (57, 58) (this comparison is reasonable assuming that *Bdellovibrio* in total can prey on about half of all bacteria that phage in total can prey on). This is despite the fact that phage have several disadvantages that we did not consider. First, half of the sequenced bacterial genomes contain a CRISPR-CAS system providing adaptive immunity against phage (59); restriction enzymes cutting up phage DNA are also common (60). Second, bacteria can rapidly evolve resistance to phages (61), in contrast to *Bdellovibrio*, although phenotypic plasticity of prey can afford temporary protection against *Bdellovibrio* (18, 44). Third, phage require a metabolically active host cell to replicate whereas *Bdellovibrio* can consume dormant prey (4). Fourth, phage do not benefit from chemotaxis as *Bdellovibrio* likely does. Nevertheless, phage outnumber *Bdellovibrio*, suggesting that these four factors can be overcome by the huge diversity of phage.

There are several factors which may promote the survival of BALOs in the natural environment. Firstly, natural environments are spatially heterogeneous, facilitating spatial refuges promoting the survival of prey species and by extension the survival of BALOs (19, 20, 62). Secondly, characteristics of the environment can make predation less efficient (giving prey a greater opportunity to recover) (63–66). Thirdly, seasonal changes can result in a temporal refuge, with prey concentrations varying with temperature (67, 68). Other temporal fluctuations could create hotspots and refuges. Fourthly, other predators, such as protists, can stabilize prey populations as they can prey on BALOs (69) and can also permit the survival of prey species in a mixed microbial community (70). Finally, prey diversity itself promotes the survival of a generalist predator, such as *Bdellovibrio*, and the presence of *Bdellovibrio* has been shown to positively correlate with alpha-diversity (68, 71–73). Overall, our results suggest that *Bdellovibrio* must prey on several prey species and locate transient hotspots to survive. Indeed, all known *Bdellovibrio* and like organisms have a wide prey range (4, 53, 74, 75).

One might expect that prey cells have to be large enough to give rise to two predator offspring, although it has been reported that it is possible for *Bdellovibrio* to start replication within one prey cell and complete this in a second prey (76). Surprisingly, our model predicts that, in all conditions tested, *Bdellovibrio* needs prey that is at least 7 times larger than itself. Indeed, this is about the difference in size between *Bdellovibrio* and *E. coli* (21) and other typical prey are of similar size. However, the predicted optimal prey size is considerably larger. Maybe *Bdellovibrio* cannot be much smaller than it already is, or accessing diverse prey species avoids precarious oscillations.

Our model also predicts that a predator that kills its prey too efficiently (has a too high attack rate, or too little mortality) will drive its prey to extinction and become extinct itself, a tragedy of the commons similar to over-fishing (36). Indeed, *Bdellovibrio* has a much higher mortality than other bacteria. While this may make over-exploitation of prey less likely, the high mortality is likely caused by its high energy expenditure when swimming fast and not feeding at the same time, also its small size prevents storage of energy reserves (6).

In conclusion, our model results suggest that *Bdellovibrio* and like organisms are unlikely to survive in most natural environments if they were preying only on a single prey species. They would also be outcompeted by phage. In line with empirical evidence, *Bdellovibrio* ought to be a generalist predator and would only thrive in prey density hotspots – which it should be able to find by chemotaxis. For application as a living antibiotic to reduce the abundance of pathogens or antimicrobial resistant bacteria in aquaculture or plant and animal agriculture, *Bdellovibrio* would be expected to be more effective where multiple prey species, not only the target species, are naturally available or added artificially.

## Supporting information

All supplementary material

## Acknowledgments

We would like to thank Andrew Lovering (University of Birmingham) and Liz Sockett (University of Nottingham) for helpful discussions of *Bdellovibrio* biology and Jamie Wood (University of York) for suggestions on predator-prey modelling. KS was supported by a Biotechnology and Biological Sciences Research Council (BBSRC) UK funded, Midlands Integrative Biosciences Training Partnership (MIBTP) PhD studentship.

