## Supplementary material for "Predation strategies of the bacterium *Bdellovibrio bacteriovorus* result in overexploitation and bottlenecks": All supplementary material

ORCIDs:
Jan-Ulrich Kreft: 0000-0002-2351-224X
Kimberley Summers: 0000-0002-0092-7848

***Corresponding author**

J. Kimberley Summers

#### Model development and simulation methods

##### ***Previous models compared with our models***

The first mathematical model of predator-prey interactions was developed separately by Lotka (1) and Volterra (2):

$$\frac{dN}{dt} = \mu N - \alpha NP \qquad (1a)$$

$$\frac{dP}{dt} = \beta NP - \lambda P \qquad (1b)$$

Where $N$ is the prey and $P$ the predator. The specific growth rate of the prey is $\mu$ and the predator has a mortality rate $\lambda$. $\alpha$ and $\beta$ are constants that reflect the amount of prey consumed in order to produce a certain number of predators. This model demonstrated that predator-prey interactions could result in oscillations of population densities without any external forcing, but these oscillations (which are neutral cycles) are not robust, as changes to initial conditions, or subsequent perturbations cause the oscillations to have a different period and amplitude.

These initial models assumed that prey growth is exponential (unlimited by resources) and that predation rate is proportional to predator and prey population sizes and does not saturate. More realistic models allow for saturation of responses. Monod (3) found that the specific growth rate of bacteria is proportional to substrate concentration until it saturates at high concentrations, analogous to Michaelis-Menten enzyme kinetics:

$$\mu = \frac{\mu_{max} S}{S + K_S} \qquad (2)$$

Where $\mu_{max}$ is the maximum specific growth rate of the prey and $K_S$ the level of substrate needed for half maximum growth rate. Holling (4) introduced a mathematically identical function to describe saturation of predation rate, known as type II functional response.

Only a few microbial predator-prey studies have modelled predatory bacteria. These models vary in prey growth function, predator functional response and the presence or absence of an explicit bdelloplast stage or other delay between prey death and the birth of new predators (Table S1).

**Table S1** Previous models of *Bdellovibrio* and other relevant microbial predators, compared to our principal Model 6.

| Bdelloplast stage? | Predator Mortality? | Prey Growth | Functional response type | Batch or chemostat | Notes | Source |
|---|---|---|---|---|---|---|
| No | No | Exponential | Non-saturating | Batch | Lotka-Volterra model | (5) |
| No | Yes | Monod | Non-saturating | Chemostat | Delay between predation and birth of predators | (6) Model 1 |
| No | Yes | Monod | Non-saturating | Chemostat | As Crowley Model 1, but also includes bdellophage (phage that infect *Bdellovibrio*) | (6) Model 2 |
| No | Yes | Monod | Holling type II | Chemostat | Protist predation | (7) |
| No | Yes | Monod | Holling type II | Chemostat | | (8) Model 1 |
| Yes | No | Monod | Non-saturating | Chemostat | As Wilkinson Model 1, but also contains decoys | (8) Model 2 |
| Yes | Yes | Exponential | Non-saturating | Batch | Contains decoys and nutrient recycling | (9) |
| Yes | Yes | Monod | Non-saturating | Batch | Includes effects of serum | (10) |
| Yes | No | Exponential | Non-saturating | Batch | Gaussian function for bdelloplast maturation time | (11) |
| Yes | Yes | Monod | Holling type II | Batch | Family of models with various ingredients examined | (12) |
| Yes | Yes | Monod | Holling type II | Chemostat | Family of models with various ingredients examined | Model 6 of this study |

***Models 1 to 6***

Several versions of our model were run on the same set of test conditions to determine the effects of each part of the model (Table S2). In Model 1, there was no predator mortality and no bdelloplast stage. Model 2 was like Model 1 but used delay differential equations (DDE) to implicitly model the delay between prey death and the birth of new predators. Model 3 was like Model 1 but had predator mortality. Models 4 to 6 had both mortality and a bdelloplast stage. In Model 4, the predator had a non-saturating functional response whereas in Models 5 and 6, a Holling type II functional response was implemented. Unlike all the other models, Model 5 replaces substrate dynamics with constant inflow of prey, as if the prey were grown in a separate chemostat feeding into the chemostat with the predator.

**Table S2** Differences between model variants.

| Model | Substrate | | Prey | | | Predator | | | | Bdelloplast | | |
|---|---|---|---|---|---|---|---|---|---|---|---|---|
| | Dilution | Prey Growth | Dilution | Growth | Predation | Dilution | Predation | Bdelloplast maturation | Mortality | Dilution | Predation | Maturation |
| 1 | $(S_0-S)D$ | $\frac{-SN\mu_N}{(K_{S,N}+S)Y_{N/S}}$ | $-ND$ | $\frac{SN\mu_N}{K_{S,N}+S}$ | $\frac{-NP\mu_P}{(K_{N,P}+N)Y_{P/N}}$ | $-PD$ | $\frac{NP\mu_P}{K_{N,P}+N}$ | | | | | |
| 2 | $(S_0-S)D$ | $\frac{-SN\mu_N}{(K_{S,N}+S)Y_{N/S}}$ | $-ND$ | $\frac{SN\mu_N}{K_{S,N}+S}$ | $\frac{-NP\mu_P}{(K_{N,P}+N)Y_{P/N}}$ | $-PD$ | $\frac{NP\mu_P}{K_{N,P}+N}$ | Implicit via DDE | | | | |
| 3 | $(S_0-S)D$ | $\frac{-SN\mu_N}{(K_{S,N}+S)Y_{N/S}}$ | $-ND$ | $\frac{SN\mu_N}{K_{S,N}+S}$ | $\frac{-NP\mu_P}{(K_{N,P}+N)Y_{P/N}}$ | $-PD$ | $\frac{NP\mu_P}{K_{N,P}+N}$ | | $-mP$ | | | |
| 4 | $(S_0-S)D$ | $\frac{-SN\mu_N}{(K_{S,N}+S)Y_{N/S}}$ | $-ND$ | $\frac{SN\mu_N}{K_{S,N}+S}$ | $\frac{-NP\mu_P}{Y_{B/N}}$ | $-PD$ | $\frac{-NP\mu_P}{Y_{B/P}}$ | $k_PB$ | $-mP$ | $-BD$ | $NP\mu_P$ | $\frac{-k_PB}{Y_{P/B}}$ |
| 5 | | | $(N_0-N)D$ | | $\frac{-NP\mu_P}{(K_{N,P}+N)Y_{B/N}}$ | $-PD$ | $\frac{-NP\mu_P}{(K_{N,P}+N)Y_{B/P}}$ | $k_PB$ | $-mP$ | $-BD$ | $\frac{NP\mu_P}{K_{N,P}+N}$ | $\frac{-k_PB}{Y_{P/B}}$ |
| 6 | $(S_0-S)D$ | $\frac{-SN\mu_N}{(K_{S,N}+S)Y_{N/S}}$ | $-ND$ | $\frac{SN\mu_N}{K_{S,N}+S}$ | $\frac{-NP\mu_P}{(K_{N,P}+N)Y_{B/N}}$ | $-PD$ | $\frac{-NP\mu_P}{(K_{N,P}+N)Y_{B/P}}$ | $k_PB$ | $-mP$ | $-BD$ | $\frac{NP\mu_P}{K_{N,P}+N}$ | $\frac{-k_PB}{Y_{P/B}}$ |

##### *Model 6 description*

The differential equations developed to track the concentrations of each variable are set out below.

$$\frac{dS}{dt} = (S_0 - S)D - N\frac{\mu_N S}{(K_{S,N}+S)Y_{N/S}} \quad (3a)$$

$$\frac{dN}{dt} = N\frac{\mu_N S}{K_{S,N}+S} - ND - P\frac{\mu_P N}{(K_{N,P}+N)Y_{B/N}} \quad (3b)$$

$$\frac{dP}{dt} = k_P B - (D+m)P - P\frac{\mu_P N}{(K_{N,P}+N)Y_{B/P}} \quad (3c)$$

$$\frac{dB}{dt} = P\frac{\mu_P N}{K_{N,P}+N} - BD - \frac{k_P B}{Y_{P/B}} \quad (3d)$$

$S_0$ is the concentration of substrate flowing into the system and $D$ is the dilution rate of the system. The maximum growth rate of the prey is $\mu_N$ and its substrate half-saturation constant is $K_{S,N}$. The attack rate constant of the predator is $\mu_P$ and its prey half-saturation constant is $K_{N,P}$. For ease of reading these half-saturation constants will be referred hereafter as the prey $K$-value and the predator $K$-value. The rate of maturation of bdelloplasts into new *Bdellovibrio* is $k_P$ and the mortality of *Bdellovibrio* is $m$. All yields are expressed in the form $Y_{A/B}$, which is the yield of consumer A per resource B consumed. The $K$-value of consumer B for resource A is expressed as $K_{A,B}$.

***Remarks on yields and biomass versus particle based unit systems***

In most animal predator-prey models, the yield terms would be expressed as the gain of $Y$ predators at the expense of 1 prey (where $Y$ is likely to be less than 1). The model developed here however has to be different to account for the unusual biology, in that both predator and prey are ‘consumed’ to produce a bdelloplast (which will later mature into new predators). This combined prey-predator entity does not exist in canonical predator-prey models and as is shown here, requires special treatment. If just a single yield term were to be associated with the bdelloplast resulting from predation, then identical amounts of predator and prey would have to go into making up that bdelloplast, which is not correct. Placing two yield terms, one on each producer species (prey and predator), by contrast, allows the ratios of producers going into the bdelloplast to be varied. This is also necessary for varying the relative sizes of prey and predator. Thus, a single yield term on the ‘product’ species needs to be replaced by two separate yield terms on the two ‘reactant’ species to use a chemical analogy.

The standard approach in animal predator-prey models is to track species in terms of individuals and not to attempt to balance biomass as this is not tractable in the wild. In contrast, the approach of studies of microbial growth and physiology in the laboratory is based on substrate and product concentrations rather than individual molecules, and biomass is often more conveniently determined than cell numbers. Moreover, for understanding metabolic pathways and energy metabolism as well as process modelling in biotechnology, the mass balance is an essential tool and growth and product yields are of primary interest.

These two traditions collide when modelling the predator-prey dynamics of a bacterial prey and a bacterial predator, suggesting the use of either particle-based or biomass-based unit systems. In this study, the biomass-based system is used predominantly, but for modelling phage, the particle-based system was found to be more suitable, so both systems were compared for a subset of the results and found to be in agreement. When tracking numbers of individuals of a species no attempt is made to explicitly balance biomass, as one prey cell combines with one predator cell to give one bdelloplast, so the two yields of bdelloplast per prey and bdelloplast per predator are both unity ($Y_{B/N} = Y_{B/P} = 1$). When the units are in terms of biomass however, mass is tracked explicitly, so care must be taken to ensure that the equations satisfy conservation of mass of prey and predator when forming a bdelloplast. For this it is required that the mass of the bdelloplast not exceed the combined mass of the predator cell and prey cell from which it is formed. For simplicity it is assumed that the conversion of predator and prey into a bdelloplast occurs without loss, such that the mass of the bdelloplast equals the combined mass of the predator cell and the prey cell. Any loss is best accounted for in the following conversion of bdelloplast into predator offspring, as this is more tractable experimentally. To enforce this mass balance constraint, it is necessary (for mass-based units) that the sum of the inverse yields is one (the inverse of the yield of bdelloplast mass formed per predator mass is the 'yield' of predator mass consumed per bdelloplast mass formed, analogously for the prey):

$$\frac{1}{Y_{B/P}} + \frac{1}{Y_{B/N}} = 1 \qquad (4a)$$

Hence

$$Y_{B/N} = \frac{Y_{B/P}}{Y_{B/P} - 1} \qquad (4b)$$

##### *Model Parameters*

Details of the model parameters are shown in Table 1. Parameters for the substrate and prey interactions of the model were based on *E. coli* as the prey species, growing on glucose as the substrate. The kinetics of this are relatively well studied and understood and the parameters were previously compiled from the literature (13).

For the predator, certain information is also well known. It takes 2-4 hours for bdelloplasts to mature (14) and release between 4 and 6 new predators when formed from *E. coli* prey (15). The relative sizes of the bacteria are also known and were used to calculate the yields. Predator dry mass per cell was assumed to be 1/7$^{th}$ that of the prey based on their relative sizes (15, 16). When one predator and one prey cell combined to give a bdelloplast each g of dry mass formed was at the expense of 0.125 g dry mass of predator and 0.875 g dry mass of prey. So, 1 g dry mass of prey gave 1.143 g dry mass of bdelloplast ($Y_{B/N}$) and 1 g dry mass of predator gave 8 g dry mass of bdelloplast ($Y_{B/P}$). For the yield of predators from bdelloplasts ($Y_{P/B}$), one bdelloplast gave 3.5 predators (15), each 1/8$^{th}$ the size, since a bdelloplast contained the mass of both the predator and the prey that formed it. Hence 1 g dry mass of bdelloplast gave 3.5/8 = 0.438 g dry mass of predator. Mortality rates were based on observations by Hespell et al. (17).

##### *Model Simulation*

The model was run in MatLab 8.6.0.267246 (R2015b) using the Ode45 solver, which implements an explicit Runge-Kutta method (18). None of the other ODE solvers were robust and accurate enough. Also, the relative and absolute tolerances of Ode45 had to be set to the very low values of 1 x $10^{-9}$. Additionally, all variables were forced to be non-negative, and all other options left as default. After every 100 hours of simulated time (equivalent to 5 volume changes at a dilution rate of 0.05 $h^{-1}$), it was checked whether the system had reached a steady state, was in sustained oscillations, or had still to reach its final state. The system was considered to be in a steady state if the maximum and minimum values of each variable were within a (settable) relative tolerance of each other for a (settable) number of hours. The system was considered to be in sustained oscillations if two consecutive peaks in the substrate values were within a (settable) relative tolerance of each other. If the system was neither in a steady state nor in sustained oscillations after 8,000 hours (equivalent to 400 chemostat volume changes at a dilution rate of 0.05 $h^{-1}$), it was deemed unstable.

##### *Parameter Sweeping*

Parameter sweeps were performed for both simulations and analytical calculations. For the simulation sweeps, the final concentration of each variable was plotted, unless the system displayed sustained oscillations, when instead the average over one oscillatory cycle was used. This average was calculated over a period between two consecutive peaks and was obtained by averaging all values recorded during this period, weighted by the time step size between recordings. For analytical sweeps the calculated steady state values for the parameter settings were plotted, as were the eigenvalues of the system and the type of regime predicted, e.g., damped oscillations. Sweeps were made in small increments through various parameters. Other parameters were set according to Table 1.

##### *Global Sensitivity Analysis*

A global sensitivity analysis was carried out on all seven parameters obtained by dimensional analysis (see SI Results). To avoid bias in the analysis, we used a Latin hypercube design (19) to obtain 10,000 tuples of parameters that were both randomly chosen and well distributed in each dimension, using the MatLab function lhsdesign to create five sets of tuples and select the best set by maximising the minimum distance between any two points to reduce clumping of tuples in parameter space. Each of these tuples was scaled evenly across the realistic parameter range (see Table 2 for minimum and maximum values), to give a set of parameter values (parameter value = minimum value + design point value * parameter range). For each set of parameters created, the steady state values were calculated using equations 9a-d. Each of the parameters was varied 1% up and down. The percentage difference between the perturbed values and the baseline values was stored for further analysis.

***Competitions***

Competing two species following different strategies is an unbiased and unequivocal way to compare the fitness of their strategies. This meant that we could compare effects, such as changes in rates at which *Bdellovibrio* consumes prey, at the expense of its economy of prey resource use. Competitions also allow a comparison of the effectiveness of *Bdellovibrio* versus bacteriophage as alternatives to antibiotics. To enable competitions, a second predator species with a second bdelloplast stage was introduced. Equation (3a) for the substrate was unaffected. Equation (3b) for the prey species became:

$$\frac{dN}{dt} = N\frac{\mu_N S}{K_{S,N}+S} - ND - P_1\frac{\mu_{P_1} N}{(K_{N,P_1}+N)Y_{B1/N}} - P_2\frac{\mu_{P_2} N}{(K_{N,P_2}+N)Y_{B2/N}} \qquad (5)$$

Equations (3c & 3d) were duplicated for $P_2$ and $B_2$, the second predator and its bdelloplast or infected cell stage.

***High Rate Predator Adjustments***

For the simulations of high rate or high yield predators, the standard settings were used for the high yield predator. The high rate predator was assumed to lyse the bdelloplast before all the available nutrients were consumed in order to more quickly release its offspring into the environment, to locate and prey upon more prey cells. As a consequence, the yield of high rate predators per bdelloplast ($Y_{P/B}$) was halved to 0.313 mg predator mg bdelloplast$^{-1}$. This reduction in yield increased the rate at which new predator biomass was formed as this rate is dependent upon the yield. To further reflect the faster maturation rate of high rate bdelloplasts, the $k_p$ was increased by a third to 0.278 mg predator mg bdelloplast$^{-1}$ h$^{-1}$. This higher rate of predator growth has not been observed in *Bdellovibrio*. However, chemostats generally select for faster growth so long-term growth of predators in a chemostat might therefore be expected to select for faster growing predators.

##### ***Bacteriophage models***

For modelling bacteriophage predation, the 'bdelloplast' stage represents the phage infected cell stage. Both in *Bdellovibrio* and phage predation, there is a lag between consumer entry and release of offspring, during which the prey cell does not grow.

Attempts to numerically solve the model using a biomass-based unit system with parameters appropriate for a bacteriophage were unsuccessful. All the ODE solvers tried were unable to resolve the rate equations in a reasonable time period. All bacteriophage simulations were therefore run using a particle-based unit system. When converting between mg dry biomass and unit cells, a prey cell is assumed to be 1 fl in size (16). The cell is assumed to be 20% dry biomass, giving 0.2 pg dry mass per unit cell. Predator dry mass per cell is assumed to be 1/7$^{th}$ that of the prey based on their relative sizes (15, 16).

In this unit system, one bdelloplast, or one infected cell, is formed from one predator cell, or one virion, and one prey cell ($Y_{B/P} = Y_{B/N} = 1$). One bdelloplast is assumed to yield 3.5 *Bdellovibrio* (14), so $Y_{P/B} = 3.5$ and one cell infected by a T4 virulent bacteriophage of *E. coli* is assumed to give 64 phage virions ($Y_{P/B} = 64$) [ref. 19].

The lysis time for T4 phage infected cells is assumed to be 36 minutes (20), giving a rate of virion production ($k_P$) of 107 hour$^{-1}$. Data for T4 from Stent & Wollman (21) were used to calculate an attack rate constant ($\mu_P$) of 19.3 infected cells predator$^{-1}$ h$^{-1}$ and a predator $K$-value ($K_{N,P}$) of 1.32 x 10$^{8}$ prey cells ml$^{-1}$.

### Results

***Model implementation and validation***

**Solver choice**. Results from ODE model simulations can be affected by the choice of ODE solver. Different solvers exist because the best choice of solver depends on the nature of the ODEs and parameters of the particular model being investigated. In order to find the most appropriate solver for this model, we ran simulations with a number of parameter settings (corresponding to the different dynamic regimes shown in Fig. 2) using MatLab solvers Ode15s, Ode23, Ode23s, Ode23t, Ode23tb, Ode45 and Ode113. All solvers were tested with absolute and relative tolerances set to the very low value of $1 \times 10^{-9}$, and all variables constrained to be non-negative (provided the solver permitted this). While all solvers tested could correctly handle conditions where the linear stability analysis had predicted a steady state (Fig. 2a, f), solvers Ode113, Ode15s, Ode23t and Ode23tb could not handle conditions where sustained oscillations were predicted (Fig. S1). Solvers Ode23 and Ode23s could handle all the scenarios, except for predicted oscillations when the initial predator numbers were increased from $1 \times 10^{-3}$ mg ml$^{-1}$ to $5 \times 10^{-2}$ mg ml$^{-1}$ for Ode23 or to $1 \times 10^{-1}$ mg ml$^{-1}$ for Ode23s. Under these conditions, simulations with the Ode23 solver incorrectly led to a sterile state with all bacteria eliminated (Fig. S1e). Solver Ode23s (which cannot be set to not allow negative values) blew up (it generated an exponential increase of substrate concentrations to the order of $10^{154}$ mg ml$^{-1}$ with negative density of prey in the order of $-1 \times 10^{154}$ mg ml$^{-1}$) (Fig. S1f). In both cases, solver Ode45 (Fig. S1g) gave sustained, extreme oscillations. Although we expected the solvers for stiff systems to be appropriate, we found that the non-stiff Ode45 solver was the only reliable one under all tested conditions, so it was used for all further investigations.

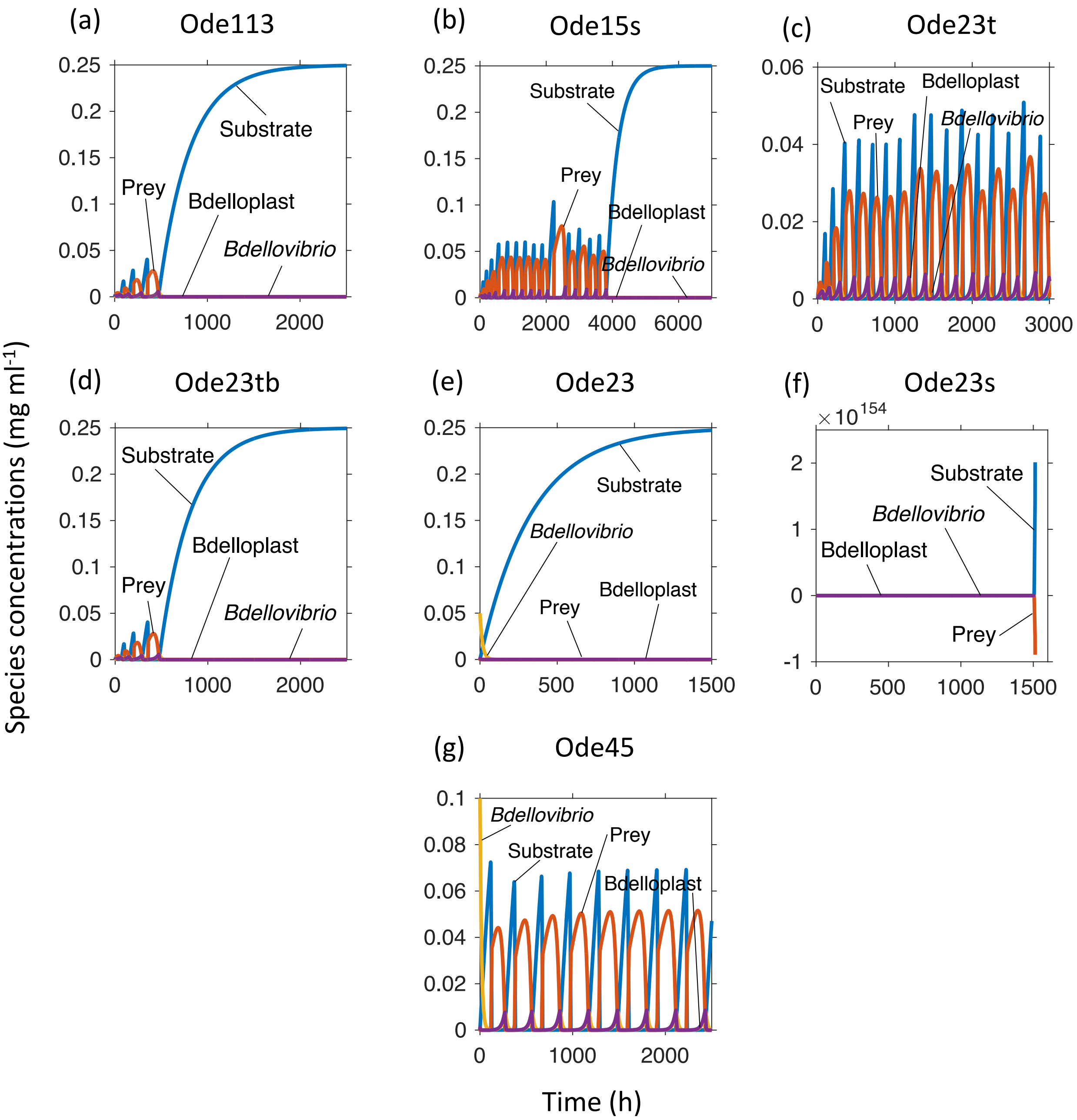

**Fig. S1** Ode45 was the only reliable and accurate solver as demonstrated by simulation results with standard parameters and $S_0$ = 0.25 mg ml$^{-1}$ and $D$ = 0.003 h$^{-1}$.

**Solver options.** We then tested a range of tolerance settings for the Ode45 solver, as tolerances that are too loose can result in inaccurate simulations, whilst overly strict tolerances are inefficient (Fig. S2). Absolute tolerance had a much stronger influence than relative tolerance. An absolute tolerance of 1 x $10^{-9}$ was needed to achieve accurate simulations. With this absolute tolerance, a relative tolerance of 1 x $10^{-6}$ or lower was needed. Reducing the relative tolerance to 1 x $10^{-9}$ produced a small improvement in the final result. Based on these finding, all further simulations were run with both absolute and relative tolerances of 1 x $10^{-9}$, unless otherwise noted.

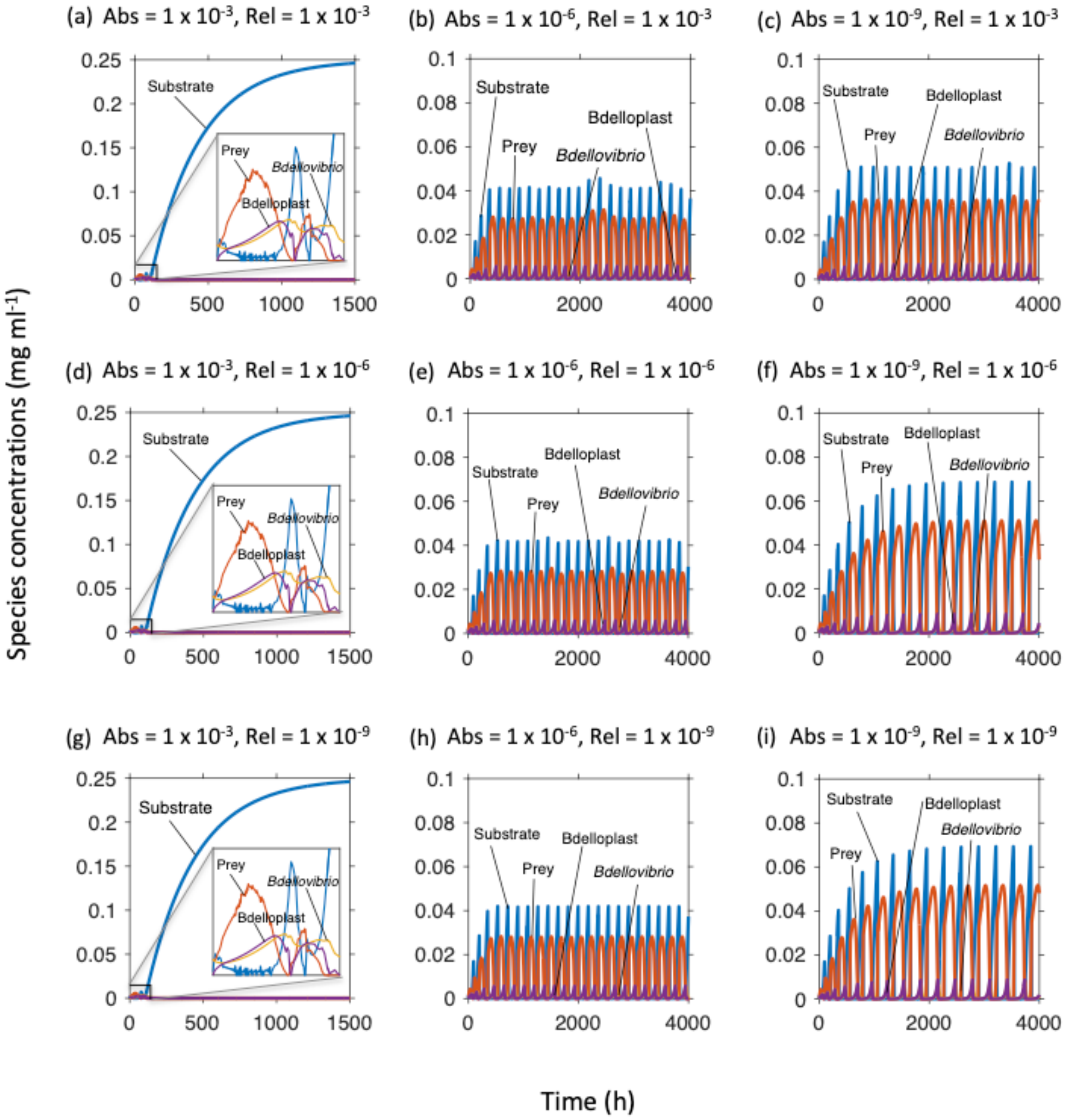

**Fig. S2** Effects of Ode45 solver tolerances on simulations with standard parameters and $S_0$ = 0.25 mg ml$^{-1}$ and $D$ = 0.003 h$^{-1}$. Absolute tolerance decreases (becomes stricter) from left to right; relative tolerance decreases from top to bottom. The strictest settings of panel **i** were used for all simulations in this study unless stated otherwise.

**Particle-based versus biomass-based units for species densities.** Most simulations were run using units of biomass density (mg dry mass ml$^{-1}$). When this was attempted with bacteriophage predators, which only differ in parameter settings from *Bdellovibrio*, simulations ran for several days without completing and eventually crashed MatLab. This was probably due to the solver being unable to find a suitable time step, as the large differences in densities of the species led to large differences in the rate of change of species over a time step. To handle these scenarios numerically, the biomass-based unit system had to be changed to a particle-based unit system (cells or particles ml$^{-1}$). This resulted in simulations completing within minutes to a few hours. We confirmed that the only effect of this change of units was to allow successful completion of simulations that would otherwise crash MatLab by rerunning simulations that could be completed with biomass-based units using particle-based units. The results from the simulations with particle-based units (Fig. S3) were equivalent to those with biomass-based units (Fig. 2), considering that the relative proportions of the species changed only apparently, since prey cells were set to be seven times larger and heavier than predator cells. There was one exception for a dilution rate of 0.0025 h$^{-1}$, where the observed pattern differed between particle- and biomass-based units. This dilution rate was very close to the boundary of a Hopf bifurcation between damped and sustained oscillations (Fig. 2a), where simulation results were very sensitive to initial conditions. For both unit systems, the initial conditions could be adjusted to obtain either dampened or sustained oscillations. For the simulation using particle-based units, the initial conditions were adjusted slightly to best match the outcome from biomass-based units.

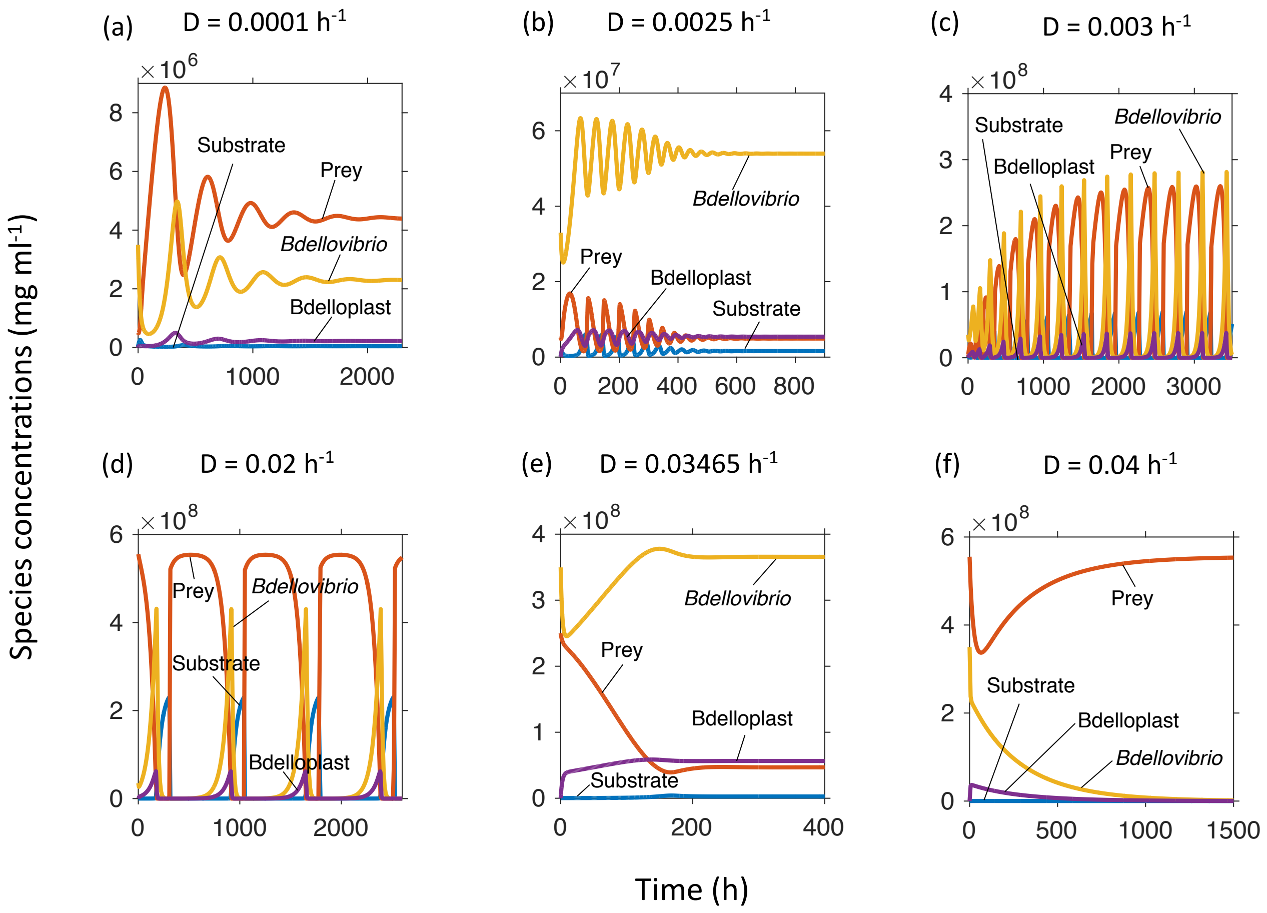

**Fig. S3** Results do not depend on the choice of unit system. Here, simulations use particle-based units, while in Fig. 2 biomass-based units were used, giving the same outcome and dynamic regimes under the same conditions ($S_0$ = 0.25 mg ml$^{-1}$, various dilution rates as in the corresponding panels in Fig. 2).

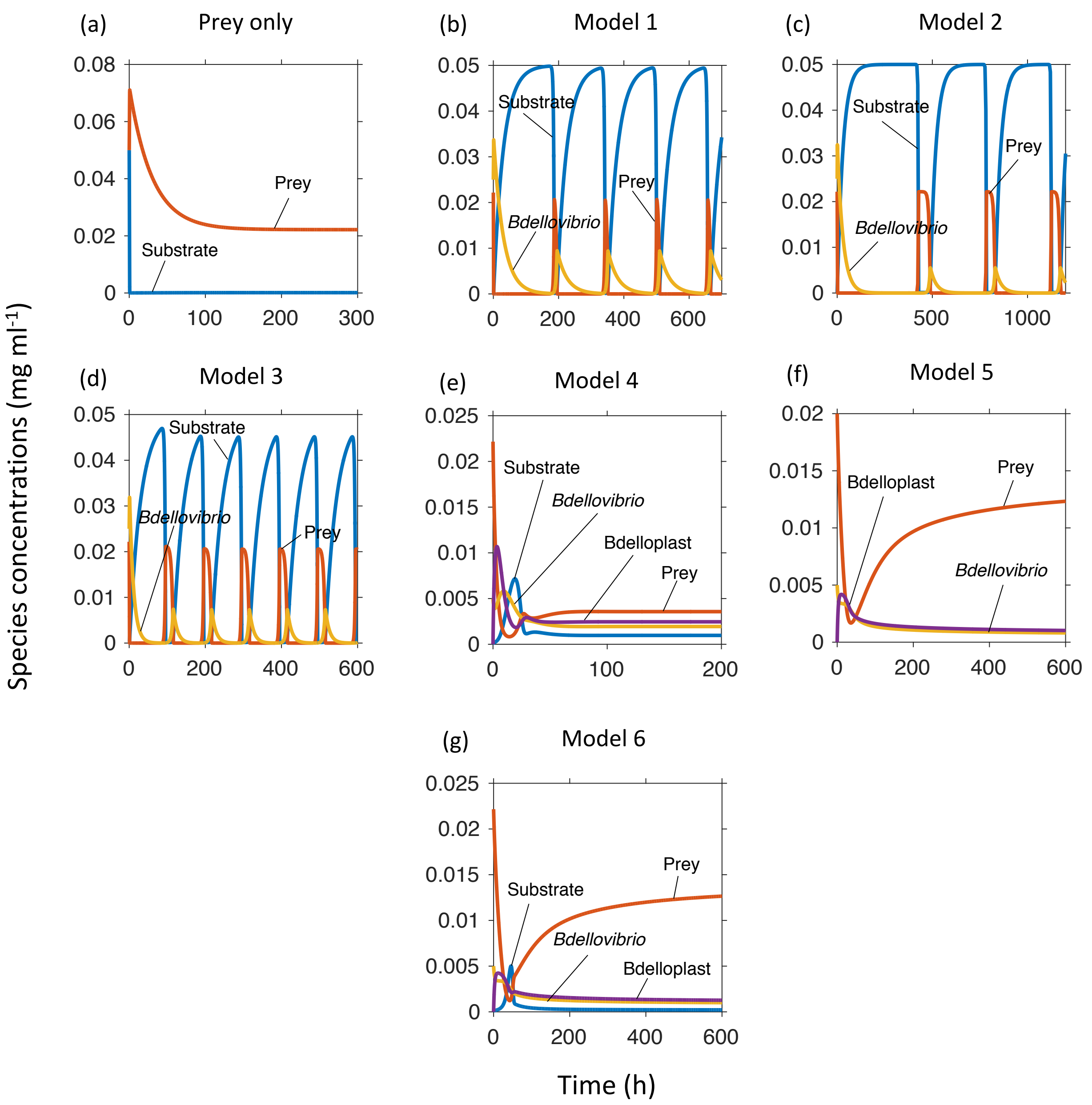

**Fig. S4** Modelling choices affect the qualitative dynamics. In all cases, the same set of default conditions, based on *Bdellovibrio* predating *E. coli* that are growing on glucose (0.05 mg ml$^{-1}$) and a dilution rate of 0.0333 h$^{-1}$ (equivalent to a 30 hour retention time) were used. This dilution rate is substantially less than 1.175 h$^{-1}$, the critical dilution rate for *E. coli* growing on this concentration of glucose in the absence of predation. All models had a Holling type II predator functional response unless otherwise stated (cf. Table S2). **a** Model 1 run without any predators (the same result was obtained for all models). **b** Model 1: without bdelloplast stage. **c** Model 2: with delay equations implicitly modelling the bdelloplast stage. **d** Model 3: as Model 1, but with predator mortality. **e** Model 4: incorporating a bdelloplast stage, predator mortality and a non-saturating functional response. **f** Model 5: as Model 4, but with constant prey input into the system instead of substrate inflow, **g** Model 6: Final model with bdelloplast stage, predator mortality and a Holling type II functional response.

##### ***Model analysis***

##### **Dimensional Analysis**

Model 6 has twelve parameters, many of which are interconnected in various ways. The more parameters a model has, the more difficult it is to determine what and how strong an effect the parameters (or their combinations) have on model outcomes. In order to reduce the number of parameters to be considered independently, and deduce any relationships between parameters, a dimensional analysis was performed. The set of dimensional equations for Model 6 were already given but are shown again for convenience:

$$\frac{dS}{dt} = (S_0 - S)D - N\frac{\mu_N S}{(K_{S,N}+S)Y_{N/S}} \tag{3a}$$

$$\frac{dN}{dt} = N\frac{\mu_N S}{K_{S,N}+S} - ND - P\frac{\mu_P N}{(K_{N,P}+N)Y_{B/N}} \tag{3b}$$

$$\frac{dP}{dt} = k_P B - (D+m)P - P\frac{\mu_P N}{(K_{N,P}+N)Y_{B/P}} \tag{3c}$$

$$\frac{dB}{dt} = P\frac{\mu_P N}{K_{N,P}+N} - BD - \frac{k_P B}{Y_{P/B}} \tag{3d}$$

First the variables were explicitly written as products of numbers and units to give:

$$\frac{d(S'S^\wedge)}{d(t'\tau)} = (S_0 - S'S^\wedge)D - N'N^\wedge\frac{\mu_N S'S^\wedge}{(K_{S,N}+S'S^\wedge)Y_{N/S}} \tag{6a}$$

$$\frac{d(N'N^\wedge)}{d(t'\tau)} = N'N^\wedge\frac{\mu_N S'S^\wedge}{K_{S,N}+S'S^\wedge} - N'N^\wedge D - P'P^\wedge\frac{\mu_P N'N^\wedge}{(K_{N,P}+N'N^\wedge)Y_{B/N}} \tag{6b}$$

$$\frac{d(P'P^\wedge)}{d(t'\tau)} = k_P B'B^\wedge - (D+m)P'P^\wedge - P'P^\wedge\frac{\mu_P N'N^\wedge}{(K_{N,P}+N'N^\wedge)Y_{B/P}} \tag{6c}$$

$$\frac{d(B'B^\wedge)}{d(t'\tau)} = P'P^\wedge\frac{\mu_P N'N^\wedge}{K_{N,P}+N'N^\wedge} - B'B^\wedge D - \frac{k_P B'B^\wedge}{Y_{P/B}} \tag{6d}$$

where primes denote numbers, carets denote units and $\tau$ is the time unit.

Next, the equations were divided by the units $S^\wedge$, $N^\wedge$, $P^\wedge$ and $B^\wedge$ and multiplied by $\tau$ to remove these from the left-hand side of the equations:

$$\frac{dS'}{dt'} = \left(\frac{S_0}{S^\wedge} - S'\right)\tau D - \tau N'N^\wedge\frac{\mu_N S'}{(K_{S,N}+S'S^\wedge)Y_{N/S}} \tag{7a}$$

$$\frac{dN'}{dt'} = N'\frac{\tau\mu_N S'S^\wedge}{(K_{S,N}+S'S^\wedge)} - N'\tau D - P'P^\wedge\frac{\tau\mu_P N'}{(K_{N,P}+N'N^\wedge)Y_{B/N}} \tag{7b}$$

$$\frac{dP'}{dt'} = \frac{\tau k_P B'B^\wedge}{P^\wedge} - (D\tau + m\tau)P' - P'\frac{\tau\mu_P N'N^\wedge}{(K_{N,P}+N'N^\wedge)Y_{B/P}} \tag{7c}$$

$$\frac{dB'}{dt'} = \frac{P'P^\wedge}{B^\wedge} \frac{\tau \mu_P N' N^\wedge}{(K_{N,P} + N' N^\wedge)} - B' \tau D - \frac{\tau k_P B'}{Y_{P/B}} \quad (7d)$$

Then we choose units to eliminate parameters ($\tau = \frac{1}{D}$, $S^\wedge = K_{S,N}$, $N^\wedge = K_{N,P}$, $P^\wedge = \frac{K_{N,P} Y_{B/N}}{Y_{B/P}}$ and $B^\wedge = K_{N,P} Y_{B/N}$) to give:

$$\frac{dS'}{dt'} = \left(\frac{S_0}{K_{S,N}} - S'\right) - N' \frac{\mu_N S' K_{N,P}}{D K_{S,N}(1+S') Y_{N/S}} \quad (8a)$$

$$\frac{dN'}{dt'} = N' \frac{\mu_N S'}{D(1+S')} - N' - P' \frac{\mu_P N'}{D(1+N') Y_{B/P}} \quad (8b)$$

$$\frac{dP'}{dt'} = \frac{k_P B' Y_{B/P}}{D} - \left(1 + \frac{m}{D}\right) P' - P' \frac{\mu_P N'}{D(1+N') Y_{B/P}} \quad (8c)$$

$$\frac{dB'}{dt'} = \frac{\mu_P N' P'}{Y_{B/P} D(1+N')} - B' - \frac{k_P B'}{D Y_{P/B}} \quad (8d)$$

Finally, we identified seven independent, dimensionless parameters of Model 6:

DP1 $\mu'_N = \frac{\mu_N}{D}$

DP2 $\mu'_P = \frac{\mu_P}{D Y_{B/P}}$

DP3 $k'_P = \frac{k_P}{D Y_{P/B}}$

DP4 $m' = \frac{m}{D}$

DP5 $S'_0 = \frac{S_0}{K_{S,N}}$

DP6 $K'_R = \frac{K_{N,P}}{K_{S,N} Y_{N/S}}$

DP7 $Y'_{B*P} = Y_{P/B} Y_{B/P}$

to give dimensionless ODEs:

$$\frac{dS}{dt} = S'_0 - S - \frac{S N \mu'_N K'_R}{1+S} \quad (9a)$$

$$\frac{dN}{dt} = \frac{S N \mu'_N}{1+S} - N - \frac{N P \mu'_P}{1+N} \quad (9b)$$

$$\frac{dP}{dt} = k'_P B Y'_{B*P} - (1 + m')P - \frac{N P \mu'_P}{1+N} \quad (9c)$$

$$\frac{dB}{dt} = \frac{N P \mu'_P}{1+N} - B - k'_P B \quad (9d)$$

Four of the seven dimensionless parameters (DP1 – DP4) are related to the dilution rate of the system as they are ratios of rates relating to the four intrinsic biological processes occurring: prey growth (DP1), predation of prey to form a bdelloplast (DP2), maturation of that bdelloplast into new predators (DP3) and predator mortality (due to starvation – DP4). The other three parameters are mass ratios. $S_0'$ (DP5) is the ratio of substrate concentration entering the chemostat to the prey $K$-value. $K_R'$ (DP6) is the ratio of predator and prey $K$-values, adjusted for the economy with which the prey converts substrate. The final parameter $Y_{B*P}'$ (DP7) is the burst size of the *Bdellovibrio*, i.e., the number of new predators formed from a single bdelloplast.

**Analytical Determination of Steady States**

Simulating the ODE system tracks population densities over time but the numerical integration can fail or be relatively slow and dependent on an appropriate choice of (sets of) initial conditions. Analytical treatment of the ODEs is an alternative that avoids these disadvantages, but results may not be valid far from steady states. We calculated all the nullclines of the ODE system. Each nullcline is the set of parameter values for which a particular differential equation equals zero (making the variable described by that equation constant in time). Where all the nullclines intersect, all of the variables are constant, so the system is in a steady state (stable or unstable). Calculating steady states was much quicker than simulation (numerical solution of the system), making it feasible to study how the steady state varied over a very large range of parameter values.

Analysis showed that for any particular set of parameters, there were up to three steady states. One only had substrate present but no organisms. A second steady state had substrate and prey, but no predators or bdelloplasts. The final potential steady state had substrate, prey and predator co-existing and as a consequence also the bdelloplast. We were particularly interested in this co-existence state. To calculate the species values at the co-existence state, equation 9d was first rearranged to describe $B$ in terms of $N$ and $P$.

$$\frac{NP\mu_P'}{1+N} - B - k_P'B = 0 \qquad (10)$$

$$B = \frac{NP\mu_P'}{(1+N)(1+k_P')} \qquad (11)$$

This value was substituted into equation 9c to give:

$$k_P' \frac{Y_{B*P}'NP\mu_P'}{(1+N)(1+k_P')} - (1+m')P - \frac{NPk_P'}{1+N} = 0 \qquad (12)$$

Which could be rearranged into:

$$N = \frac{(1+m')(1+k_P')}{k_P'\mu_P'Y_{B*P}' - \mu_P'(1+k_P') - (1+m')(1+k_P')} \qquad (13)$$

Equation 9a was used to give $S$ in terms of $N$:

$$S_0' - S - \frac{SN\mu_N' K_R'}{1+S} = 0 \quad (14)$$

$$S^2 + S(1 + N\mu_N' K_R' - S_0') - S_0' = 0 \quad (15)$$

This quadratic equation has two solutions, however, only the physically possible non-negative solution was recorded as a genuine steady state.

Finally, equation 9b was used to get $P$ in terms of $N$ and $S$:

$$\frac{SN\mu_N'}{1+S} - N - \frac{NP\mu_P'}{1+N} = 0 \quad (16)$$

$$P = \left(\frac{S\mu_N'}{1+S} - 1\right)\frac{(1+N)}{\mu_P'} \quad (17)$$

This meant the three equilibrium points or steady states of the system were, in order $S$, $N$, $P$, $B$:

Abiotic state:

$$(S_0', 0, 0, 0)$$

Predator free state (prey and substrate only):

$$\left(\frac{1}{\mu_N'-1}, \frac{S_0'(\mu_N'-1)-1}{K_R'(\mu_N'-1)}, 0, 0\right)$$

All species co-existence (note only 1 possible value for $S$ is non-negative):

$$\left(\frac{-(1 + N\mu_N' K_R' - S_0') \pm \sqrt[2]{(1 + N\mu_N' K_R' - S_0')^2 - 4S_0'}}{2}, \frac{(1+m')(1+k_P')}{k_P'\mu_P' Y_{B*P}' - \mu_P'(1+k_P') - (1+m')(1+k_P')}, \left(\frac{S\mu_N'}{1+S} - 1\right)\frac{(1+N)}{\mu_P'}, \frac{NP\mu_P'}{(1+N)(1+k_P')}\right)$$

**Jacobian Matrix**

The Jacobian matrix is the set of all first order partial derivatives of all differential equations of a system with respect to all variables, evaluated at a particular steady state (Fig. S5). It represents the linearized system of the non-linear ODEs at a given steady state and is used for linear stability analysis because linear systems can be solved. The caveat is that the linearization only approximates the system behaviour, and may only be valid near the steady state. The solution consists of sums of exponential functions of the form $\boldsymbol{x}(t) = \boldsymbol{v}e^{\lambda t}$ with the eigenvalues $\lambda$ of the Jacobian matrix in the exponents determining the time evolution of perturbations $\boldsymbol{x}(t)$ of the steady state and therefore the local stability of steady states (22).

Any complex eigenvalues indicate that the system will display some form of oscillations. If the common real part of the complex conjugate eigenvalue pair is negative, the oscillations will be damped, and the equilibrium point is a stable focus. While for a positive real part, the oscillations are sustained, and the equilibrium point is an unstable node, which may have an associated stable limit cycle. If all the eigenvalues are real and negative this indicates a stable steady state with the equilibrium point being a stable node. If all eigenvalues are real and at least one is positive, the state is unstable, and the equilibrium point is either a saddle node or an unstable node. In order to calculate these eigenvalues for various system parameters, the Jacobian matrix was determined for the dimensionless equations. The rows of the Jacobian matrix were derived for the substrate $S$, prey $N$, predator $P$ and bdelloplast $B$ rates of change in this order (Eqs. 9a-d) by partially differentiating by $S$, $N$, $P$ and $B$ in turn for each column (Fig. S5).

$$
\begin{array}{c|cccc}
 & \frac{\delta}{\delta S} & \frac{\delta}{\delta N} & \frac{\delta}{\delta P} & \frac{\delta}{\delta B} \\
\hline
\frac{dS}{dt} & -1-\frac{N\mu_N' K_R'}{(1+S)^2}, & -\frac{S\mu_N' K_R'}{1+S}, & 0, & 0, \\
\frac{dN}{dt} & \frac{N\mu_N'}{(1+S)^2}, & \frac{S\mu_N'}{1+S}-1-\frac{P\mu_P'}{(1+N)^2}, & -\frac{N\mu_P'}{1+N}, & 0 \\
\frac{dP}{dt} & 0, & -\frac{P\mu_N'}{(1+N)^2}, & -1-m'-\frac{N\mu_P'}{1+N} & k_P' Y_{B*P}' \\
\frac{dB}{dt} & 0, & \frac{P\mu_N'}{(1+N)^2}, & \frac{N\mu_P'}{1+N}, & -1-k_P'
\end{array}
\qquad (18)
$$

**Fig. S5** Jacobian matrix for Model 6 at four species co-existence steady state. Zero entries, circled in brown, indicate independence, i.e., changing a variable had no effect on the rate of change of the other variable. Entries that were always negative, circled in red, mean that an increase in one variable caused a decrease in the rate of change of the corresponding variable. Entries that were always positive, circled in blue, meant that an increase in one variable caused an increase in the rate of change of the other. Entries circled in green, could be positive or negative depending on parameter values.

Each entry in the Jacobian matrix (Eq. 18) describes the effect of a small change in one species on the rate of change of another species, when the system was initially at steady state. The values on the main diagonal, are the effects of a change in the density of a variable on its own rate of change. The eigenvalues of the Jacobian matrix are found using equation 19, where $\lambda$ is an eigenvalue, $J$ the Jacobian matrix and $I$ the identity matrix (22).

$$J - \lambda I = 0 \qquad (19)$$

This can in theory be solved analytically, but gives rise to a quartic equation that is intractable and has to be evaluated numerically. Instead taking inspiration from the graphical analysis method of Rosenzweig and MacArthur (23) we examined the $\frac{\delta}{\delta N}$ of $\frac{dN}{dt}$ entry of the Jacobian matrix (Eq. 18) circled in green in Fig. S5. This is the rate of change of the prey nullcline with respect to prey density. In a two-dimensional predator-prey system, when this is positive, the equilibrium point is unstable, and the result is stable oscillations, resulting from a stable limit cycle. When it is negative, the equilibrium point is stable. The point at which the steady state of the system goes from being a stable limit cycle, surrounding an unstable node, to damped oscillations around a stable node, is a Hopf bifurcation of the system. A four-dimensional system, such as described by Model 6 is inherently more complex than a two-dimensional system, but similar principles apply. Substituting the equilibrium value of:

$$P = \left(\frac{S\mu'_N}{1+S} - 1\right)\frac{(1+N)}{\mu'_P} \qquad (20)$$

into the equation for the $\frac{\delta}{\delta N}$ of $\frac{dN}{dt}$ entry in the Jacobian matrix (Eq. 18):

$$\frac{S\mu'_N}{1+S} - 1 - \frac{P\mu'_P}{(1+N)^2} \qquad (21)$$

Gave:

$$\left(\frac{S\mu'_N}{1+S} - 1\right) - \frac{\left(\frac{S\mu'_N}{1+S} - 1\right)}{1+N} \qquad (22)$$

Which can only be zero if either $N = 0$, which is not a point at which all four species are co-existing, as prey density is zero, or when $\frac{S\mu'_N}{1+S} - 1 = 0$. Substituting the equilibrium value for $S$

$$S = \frac{-\left(1 + N\mu'_N K'_R - S'_0\right) \pm \sqrt[2]{\left(1 + N\mu'_N K'_R - S'_0\right)^2 - 4S'_0}}{2} \qquad (23)$$

into equation 22 gave an equation that could be solved for inflow substrate concentration ($S_0$). Inputting the default parameters (Table 1) and a dilution rate of 0.02 h$^{-1}$ and solving gave a negative and thus impossible value for $S_0$ of -0.00607 mg ml$^{-1}$, indicating that there is no value of $S_0$ that, with these parameter values, would make the $\frac{\delta}{\delta N}$ of $\frac{dN}{dt}$ entry in the Jacobian matrix zero (or negative). This is not entirely surprising, as this is an unusual four species system in which predator and prey densities both decrease when predation events occur and only bdelloplast densities increase.

Fig. 2a shows that the system does in fact undergo Hopf bifurcations. Take as an example the series of points indicated by the series of white crosses in Fig. 2a, whose Jacobian matrix values and eigenvalues are listed in Table S3. The inflow substrate concentration ($S_0$) was set to 0.25 mg ml$^{-1}$ (and all other parameters to their default values). When the dilution rate ($D$) was set to 0.04 h$^{-1}$, the predator washed out of the system and only the prey remained (Fig. 2a, g). When $D$ was reduced to 0.03465 h$^{-1}$, the predator could multiply fast enough to survive within the chemostat and a four species equilibrium point occurred, the eigenvalues for which were all real and negative, indicating the equilibrium point was a stable node (Table S3 and Fig. 2a, f). When $D$ was lowered further to 0.003 h$^{-1}$, two of the eigenvalues became negative real numbers, the other two became a complex conjugate pair, with a positive real part, indicating the equilibrium point was an unstable node, with an associated stable limit cycle (Table S3 and Fig. 2a, d). Driving $D$ still lower to 0.0025 h$^{-1}$, resulted in the system passing through a Hopf bifurcation. The complex conjugate pair of eigenvalues associated with the equilibrium point of this system had crossed the imaginary axis and now had negative real parts, while the other two eigenvalues were both still negative real numbers, indicating the equilibrium point was a stable focus, with associated damped oscillations (Table S3 and Fig. 2a, c).

**Table S3** Eigenvalues and Jacobian matrix of the co-existence state for selected dilution rates. Parameters all had default values (Table 1), except $S_0$ = 0.25 mg ml$^{-1}$ and $D$ was as shown.

| Dilution rate ($D$) (h$^{-1}$) | Jacobian matrix | | | | Eigenvalues | Type of node |
|---|---|---|---|---|---|---|
| 0.0001 | -5.65x10$^3$ | -0.105x10$^3$ | 0 | 0 | -5.52<br>-3.34<br>-<br>0.0255+0.167i<br>-0.0255-0.167i | Stable focus (with damped oscillations) |
| | 6.83x10$^3$ | 63.4 | -2.40 x10$^2$ | 0 | | |
| | 0 | -62.0 | -84.1 | 8.72 x10$^2$ | | |
| | 0 | 0.0620 | 0.2402 | -2.4896 | | |
| 0.0025 | -93.4 | -93.1 | 0 | 0 | -139<br>-<br>0.0540+0.666i<br>-0.0540-0.666i<br>-19.7 | Stable focus (with damped oscillations) |
| | 112 | 59.4 | -10.1 | 0 | | |
| | 0 | -52.1 | -35.1 | 349 | | |
| | 0 | 52.1 | 10.1 | -101 | | |
| 0.003 | -62.5 | -90.7 | 0 | 0 | -117<br>7.10 + 57.1i<br>7.10 - 57.1i<br>-14.2 | Unstable node (with associated limit cycle) |
| | 74.4 | 58.6 | -8.53 | 0 | | |
| | 0 | -50.2 | -29.5 | 291 | | |
| | 0 | 50.2 | 8.53 | -84.0 | | |
| 0.02 | -1.14 | -27.5 | 0 | 0 | 24.0<br>-20.0<br>-1.04<br>0.983 | Unstable node |
| | 0.168 | 24.2 | -1.78 | 0 | | |
| | 0 | -8.04 | -5.78 | 43.6 | | |
| | 0 | 8.04 | 1.78 | -13.4 | | |
| 0.035 | -227 | -4.27 | 0 | 0 | -222<br>-12.2<br>-0.679<br>-0.439 | Stable node |
| | 274 | 4.00 | -1.31 | 0 | | |
| | 0 | -0.160 | -4.04 | 25.1 | | |
| | 0 | 0.160 | 1.31 | -8.16 | | |

**Dynamic regimes, linear stability analysis and dimensionless equations.**

There was usually a good agreement between the type of regime predicted from the eigenvalues of the Jacobian matrix (Table S4) and those observed in simulations (Fig. 2). Exceptions were at the boundaries between regimes and in the region predicted by the eigenvalues to be linearly unstable, where the simulations resulted in very long-period, extreme oscillations. The calculations based on the eigenvalues were using the dimensionless form of the ODEs, whilst the simulations were performed using dimensional equations. To check that any differences seen were not due to using dimensional or dimensionless equations, all scenarios from Fig. 2 were also simulated with dimensionless equations, producing identical results (data not shown). Hence, dimensional equations were used for all further simulations.

**Table S4** Possible dynamical outcomes and associated eigenvalues of the predator and prey co-existence state where it exists. Predator and prey coexistence implies the presence of the bdelloplast and substrate. Eigenvalues were not determined (ND) when not all variables were positive.

| **Dynamical regimes** | **Real parts of eigenvalues** | **Imaginary parts of eigenvalues** |
|---|---|---|
| Complete washout of all bacteria | N/A as four species co-existence state does not exist | |
| Prey only survival | N/A as four species co-existence state does not exist | |
| Stable co-existence of all species (stable node) | All negative | All zero |
| Co-existence of all species with damped oscillations (stable focus) | All negative | Not all zero |
| Co-existence of all species with sustained, stable oscillations (stable limit cycle) | At least two positive | Not all zero |
| Co-existence of all species, but linearly unstable (saddle node or unstable node) | At least one positive | All zero |

***Protist model***

The population dynamics generated by our model are qualitatively different from those generated by typical Lotka-Volterra models for animal populations (24). In Lotka-Volterra models, the oscillations are gentle and fairly similar to sine waves, whereas in our model the waves are very asymmetric, approach zero very closely and rise and fall abruptly. Such extreme oscillations were also seen by Curds and Bazin when they modelled protist predation (24), but their oscillations had a much shorter period. We sought to understand the reason for the differences in period length by recreating their model, then introducing parameters relevant to *Bdellovibrio* predation (Fig. S6). We found that decreasing the dilution rate, decreasing the predator $K$-value and reducing the attack rate constant all increased the period length.

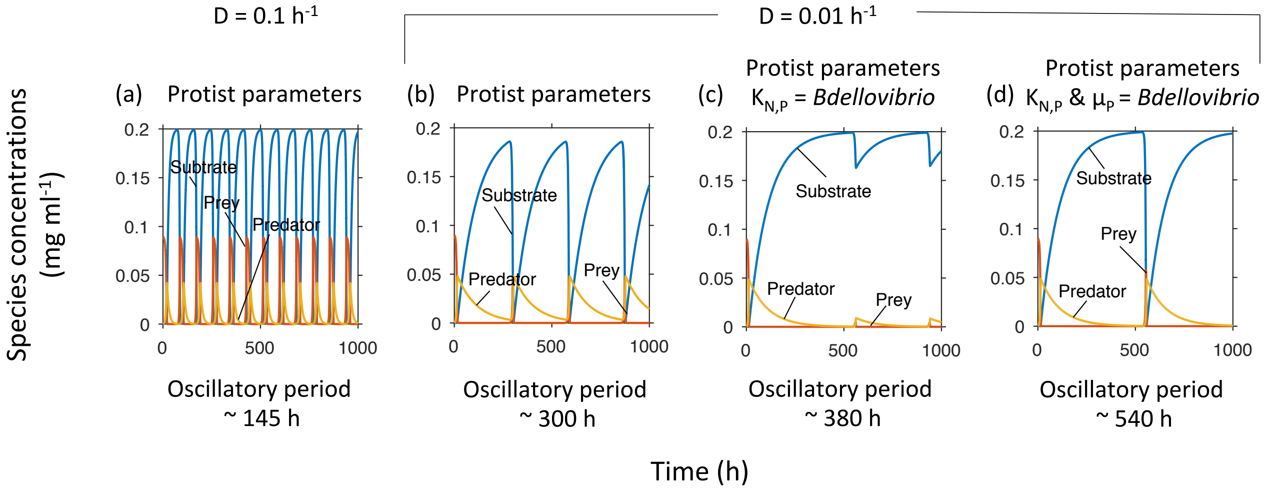

**Fig. S6** The protist model of predation (24) shows similarly extreme oscillations as the *Bdellovibrio* model but shorter periods. **a** The protist predation model without predator mortality or the equivalent to a bdelloplast stage gave a similar pattern of behaviour to that seen in our model. With a dilution rate ($D$) of 0.1 $h^{-1}$, the value used in (24), the oscillatory period was shorter than seen in our model. **b** Reducing $D$ lengthened the period. **c** Adjusting the much higher predator $K$-value of the protist ($K_{N,P}$ = 1.2 x $10^{-2}$ mg dry mass $ml^{-1}$) to that of *Bdellovibrio* ($K_{N,P}$ = 8.6 x $10^{-4}$ mg dry mass $ml^{-1}$) lengthened the oscillatory period and reduced the prey recovery, resulting in a different pattern of oscillations. **d** When the attack rate constant was also adjusted from the protist rate ($\mu_P$ = 0.43 $h^{-1}$) to that of *Bdellovibrio* ($\mu_P$ = 0.38 $h^{-1}$), the original pattern was restored but the period further lengthened.

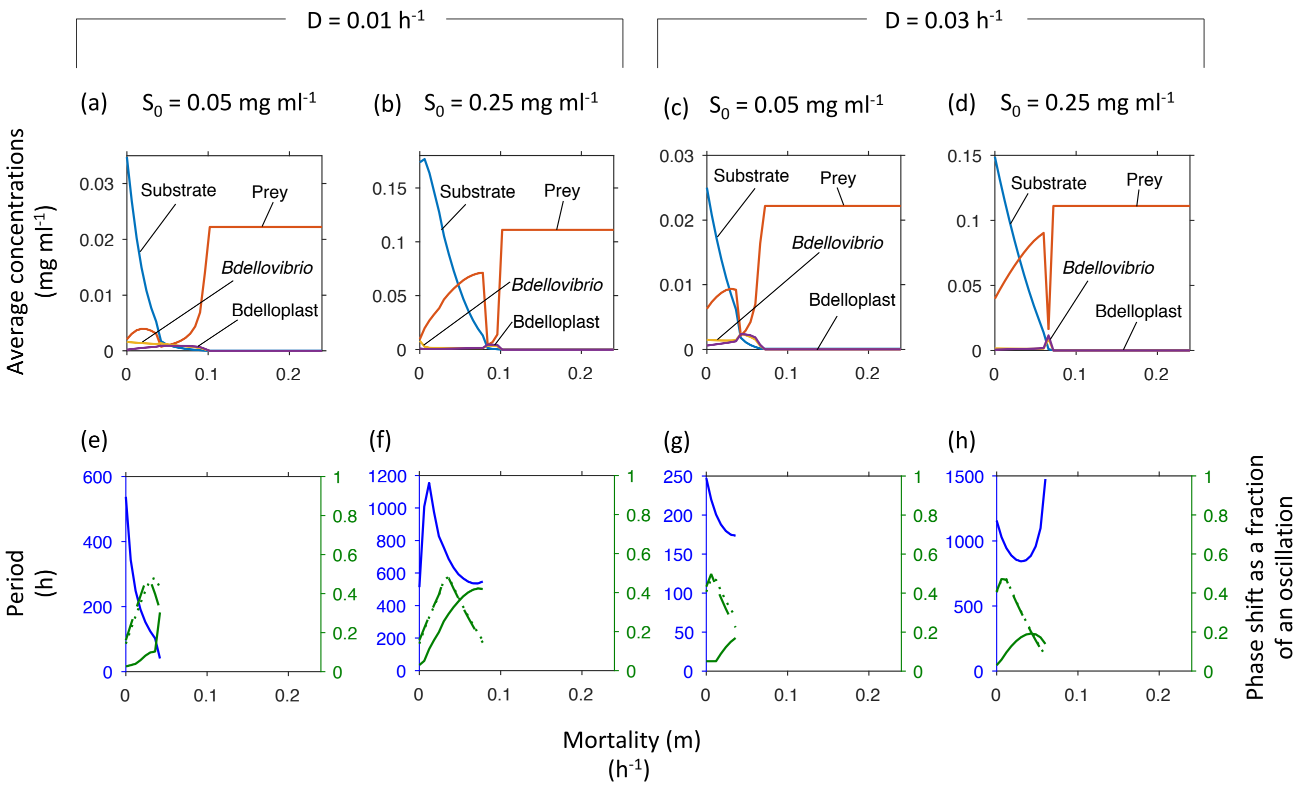

**Fig. S7** Optimal predator mortality. Above a critical predator mortality rate, the population densities no longer oscillated, prey density dropped and predator density reached a maximum, which was higher than at zero mortality. At even higher mortality, the predator died out, leading to a prey only steady state. At higher substrate inflow concentrations ($S_0$) and dilution rates, the predator abundance peak narrowed. Top row shows concentrations at steady state or averaged over one oscillatory cycle. Bottom row shows the oscillatory period (blue, left axis) and phase shifts (green, right axis) from substrate peak to peak of prey (solid line), free *Bdellovibrio* (dashed line) or bdelloplast (dotted line).

##### *Maturation rate of bdelloplast*

We swept through a range of bdelloplast maturation rates ($k_P$) from 0.02 to 0.5 mg predator mg bdelloplast$^{-1}$ h$^{-1}$ under various inflow substrate concentrations ($S_0$) and dilution rates ($D$) (Fig. S8). There was obviously a minimal $k_P$ required for predator persistence. There was also a $k_P$ that was optimal for predator density, visible at the higher $D$. This optimum was just below the $k_P$ threshold leading to oscillations at higher $S_0$ (Fig. S8). Increasing $D$ caused an increase in the minimum value of $k_P$ required for predator survival, but otherwise did not affect the trends seen (Fig. S8a, c). Increasing $S_0$ destabilised the system and resulted in oscillations (Fig S8b, d, e, f). Increasing $k_P$ generally reduced the period of oscillations where these occurred (Fig. S8e, f).

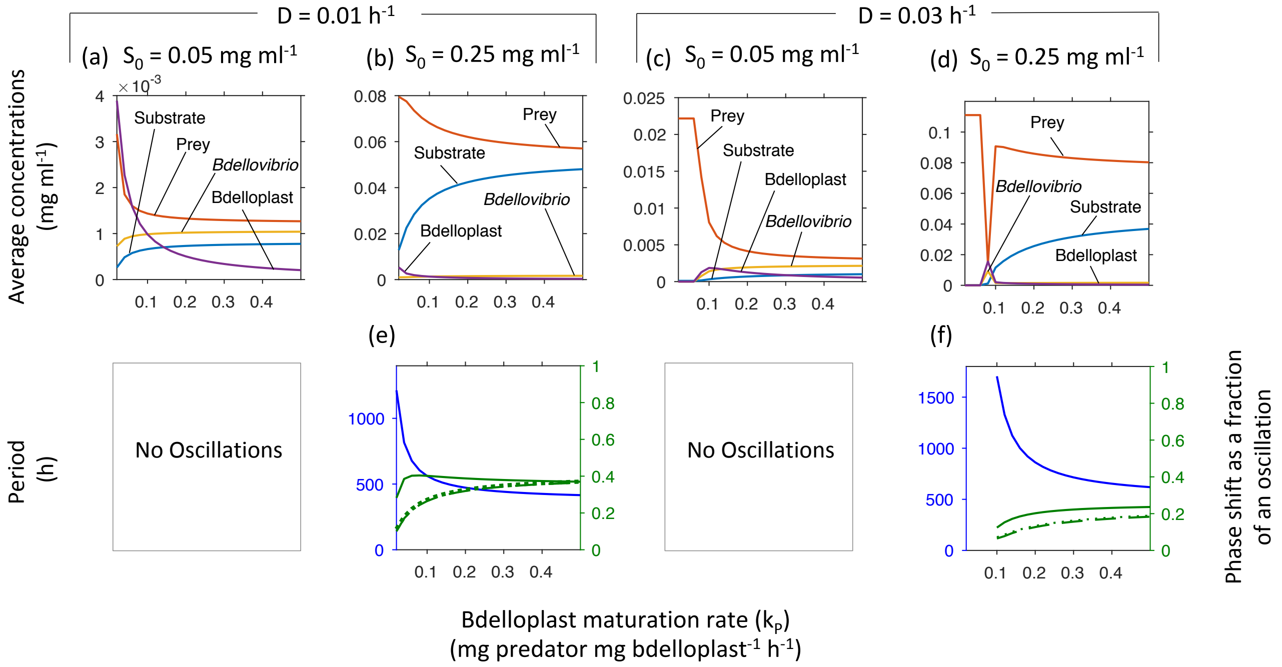

**Fig. S8** Varying the maturation rate of the bdelloplast ($k_P$) from 0.02 to 0.5 mg predator mg bdelloplast$^{-1}$ h$^{-1}$ had a plethora of effects. **a, c** At low inflow substrate concentrations ($S_0$), the populations did not oscillate regardless of maturation rate. The minimal $k_P$ for predator persistence and the optimal $k_P$ are visible in **c**. **b, d** At higher $S_0$, populations oscillated above a threshold $k_P$; the optimal $k_P$ was just below this threshold (visible in **d**). **a-d** Concentrations at steady state or averaged over one oscillatory cycle. **e, f** Oscillatory period (blue, left axis) and phase shifts (green, right axis) from substrate peak to peak of prey (solid line), free *Bdellovibrio* (dashed line) or bdelloplast (dotted line).

##### ***$K$-values of predator and prey***

One of the fundamental system parameters identified from the dimensional analysis was $K_R' = \frac{K_{N,P}}{K_{S,N} Y_{N/S}}$, the ratio of predator and prey $K$-values ($K_{N,P}$ and $K_{S,N}$), scaled by the yield of prey per substrate ($Y_{N/S}$). While $K_{S,N}$ and $Y_{N/S}$ are better understood, $K_{N,P}$ has hardly been studied. By re-analysing data from the study by Varon & Zeigler (5) of a relative of *Bdellovibrio*, the marine strain BM4, predating *Photobacterium leiognathi*, we were able to calculate an estimate for both $\mu_P$ and $K_{N,P}$. It should be noted that this involved a different predator strain predating a different prey species, and as such, this system likely has somewhat different attack kinetics. To gain an understanding of the effect of these differences we swept through a range of $K_{N,P}$ values at various inflow substrate concentrations ($S_0$). At the lowest $S_0$ values, populations did not cycle and the lowest $K_{N,P}$ was optimal (Fig. S9a). However, at higher $S_0$, low $K_{N,P}$ values resulted in oscillations with reduced average predator levels (Fig. S9b-e). Counterintuitively, raising the $K_{N,P}$ resulted in a bifurcation to a stable co-existence that benefited the predator at the expense of the prey. The optimal $K_{N,P}$ was just above this critical value. Further increases resulted in a slow decline in predator density and a sharper increase in prey, until a second threshold was reached leading to a prey only steady state (Fig. S9b). As would be expected, increasing $S_0$ increased the prey density, resulting in an increase in the $K_{N,P}$ at which the bifurcations occurred.

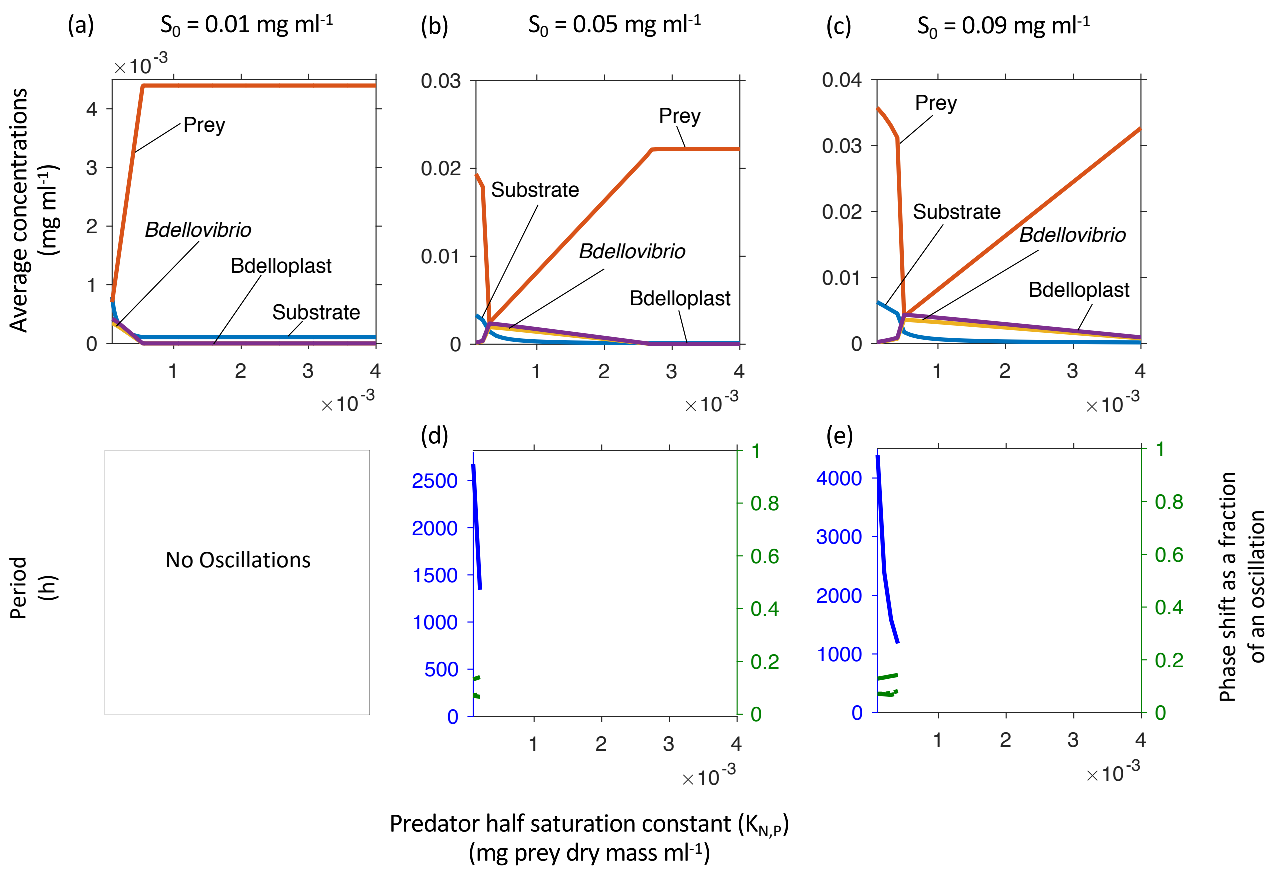

**Fig. S9** Predator $K$-value ($K_{N,P}$): lowest was not always best. **a** At the lowest inflow substrate concentration ($S_0$), the system did not oscillate and the lowest $K_{N,P}$ was best for the predator. **b-e** At higher $S_0$, a too low $K_{N,P}$ resulted in oscillations and reduced predator density. An optimal $K_{N,P}$ existed just below the critical value where oscillations stopped. Top row shows concentrations at steady state or averaged over one oscillatory cycle. Bottom row shows the oscillatory period (blue, left axis) and phase shifts (green, right axis) from substrate peak to peak of prey (solid line), free *Bdellovibrio* (dashed line) or bdelloplast (dotted line). All simulations were performed with a dilution rate of 0.03 $h^{-1}$.

***Inflow of the substrate for prey growth***

The dimensional analysis showed that, in terms of qualitative system behaviour, the inflow concentration of the substrate consumed by the prey should be seen relative to the $K_{S,N}$ ($S'_0 = S_0/K_{S,N}$). In essence, this parameter determines the productivity of the prey. It might be expected that increasing the amount of nutrients coming into a system would be of benefit to all species within that system, especially predators, which are at the top of the food chain. On the other hand, increased nutrient levels can destabilise an ecosystem, known as the paradox of enrichment effect (25). We sought to determine which of these effects would be observed in our system. As varying $K_{S,N}$ would have altered another dimensionless parameter, $K'_R$, we instead varied $S_0$. Increasing $S_0$ from 0 initially benefitted the predator more than the prey and led to a predator maximum (Fig. S10). Increasing $S_0$ beyond this led to a drop in predator abundance and a rise in prey abundance until a bifurcation occurred from stable co-existence of predator and prey to a regime of extreme oscillations. Once oscillations were seen, further increases in $S_0$ benefitted the prey but not the predator (Fig. S10a, b), while the oscillatory period increased to a maximum (Fig. S10c, d). Beyond the maximum period, and only for the lower dilution rate, further increases in $S_0$ benefitted both the predator and the prey (Fig. S10a).

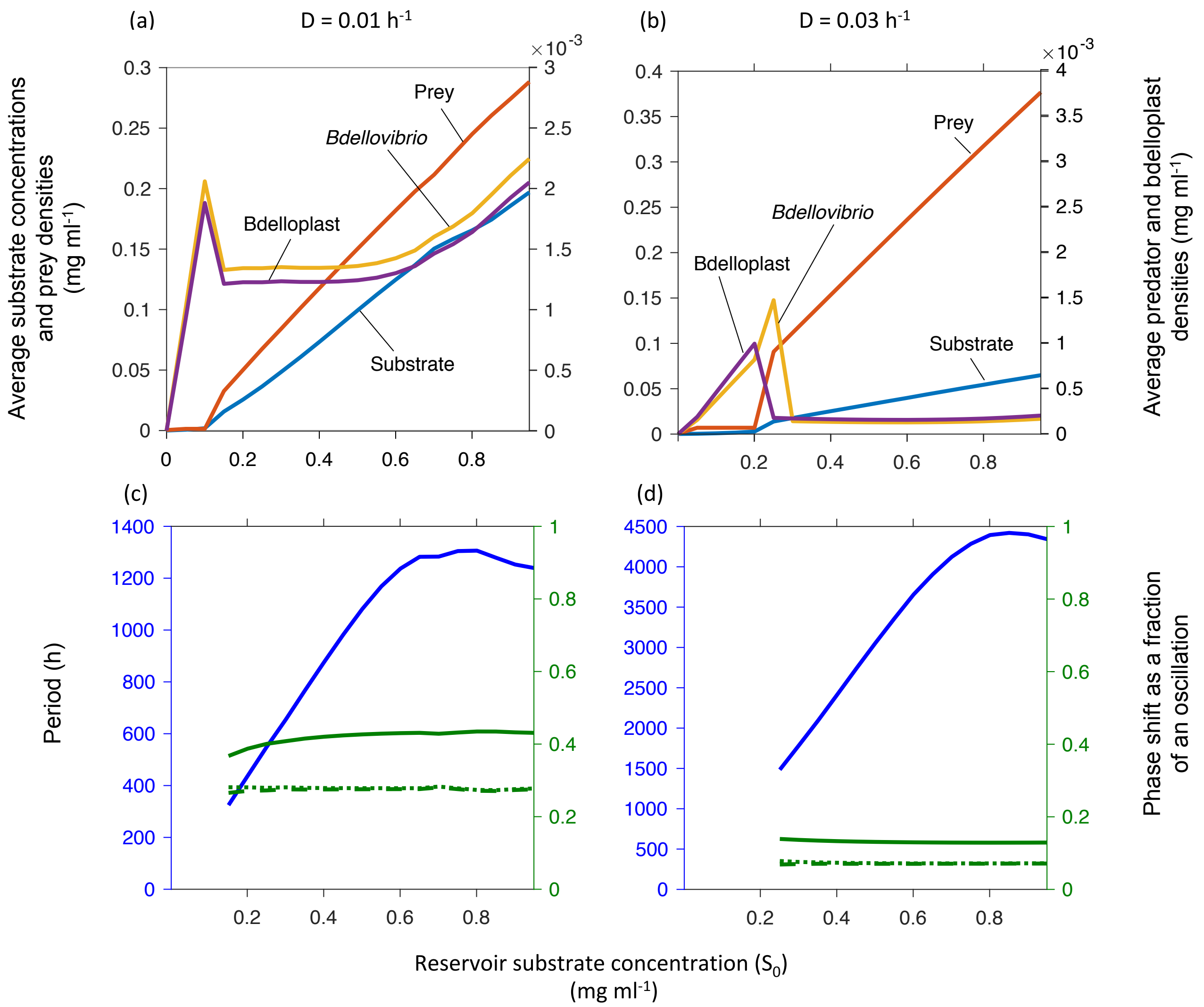

**Fig. S10** With increasing inflow substrate concentration ($S_0$), system behaviour underwent a sequence of changes. First, predator but not prey abundance increased towards a maximum and dropped thereafter. Then a bifurcation led to oscillations with increasing prey but not predator abundance and the period increased to a maximum. Only at the lower dilution rate, did the predator eventually benefit. Note predator abundances are shown on a 100-fold magnified scale. Top row shows concentrations at steady state or averaged over one oscillatory cycle. Bottom row shows the oscillatory period (blue, left axis) and phase shifts (green, right axis) from substrate peak to peak of prey (solid line), free *Bdellovibrio* (dashed line) or bdelloplast (dotted line).

***Bdelloplast burst size***

The final dimensionless parameter, $Y'_{B*P}$, corresponded to the *Bdellovibrio* burst size, i.e., the number of offspring emerging from a prey cell. Hence, $Y'_{B*P}$ increases with prey size. We hypothesised that increased burst sizes would benefit the predator as it would be using its prey more economically, whilst lower burst sizes would reflect a more wasteful use of prey. Since the two components of $Y'_{B*P}$, $Y_{B/P}$ and $Y_{P/B}$, were also components of other parameters, $\mu'_P$ and $k'_P$, respectively, altering either would change two dimensionless parameters. To prevent this, we varied both $Y_{B/P}$ and $\mu_P$ at the same time, such that their ratio was kept constant. This ensured that $Y'_{B*P}$ was varied without altering $\mu'_P$. We found a minimal and optimal burst size (Fig. S11). The lower the dilution rate, the lower the minimal and optimal burst size and the broader the optimum (Fig. S11). Increasing the burst size above the optimal value resulted in a bifurcation to extreme oscillations, corresponding to a sharp rise in prey and drop in predator average densities, but only at the low inflow substrate concentration ($S_0$). Further increases caused the densities of both predator and prey to decline (Fig. S11b, c). At higher $S_0$, the optimum was very narrow, and oscillations did not occur. Contrary to initial expectations, we found that once again, whilst a certain burst size is required for predator survival, too large a burst size, i.e. too economic a predator, results in a boom in the predator population that cannot be sustained by the prey, as it cannot reproduce quickly enough to make up for losses due to predation.

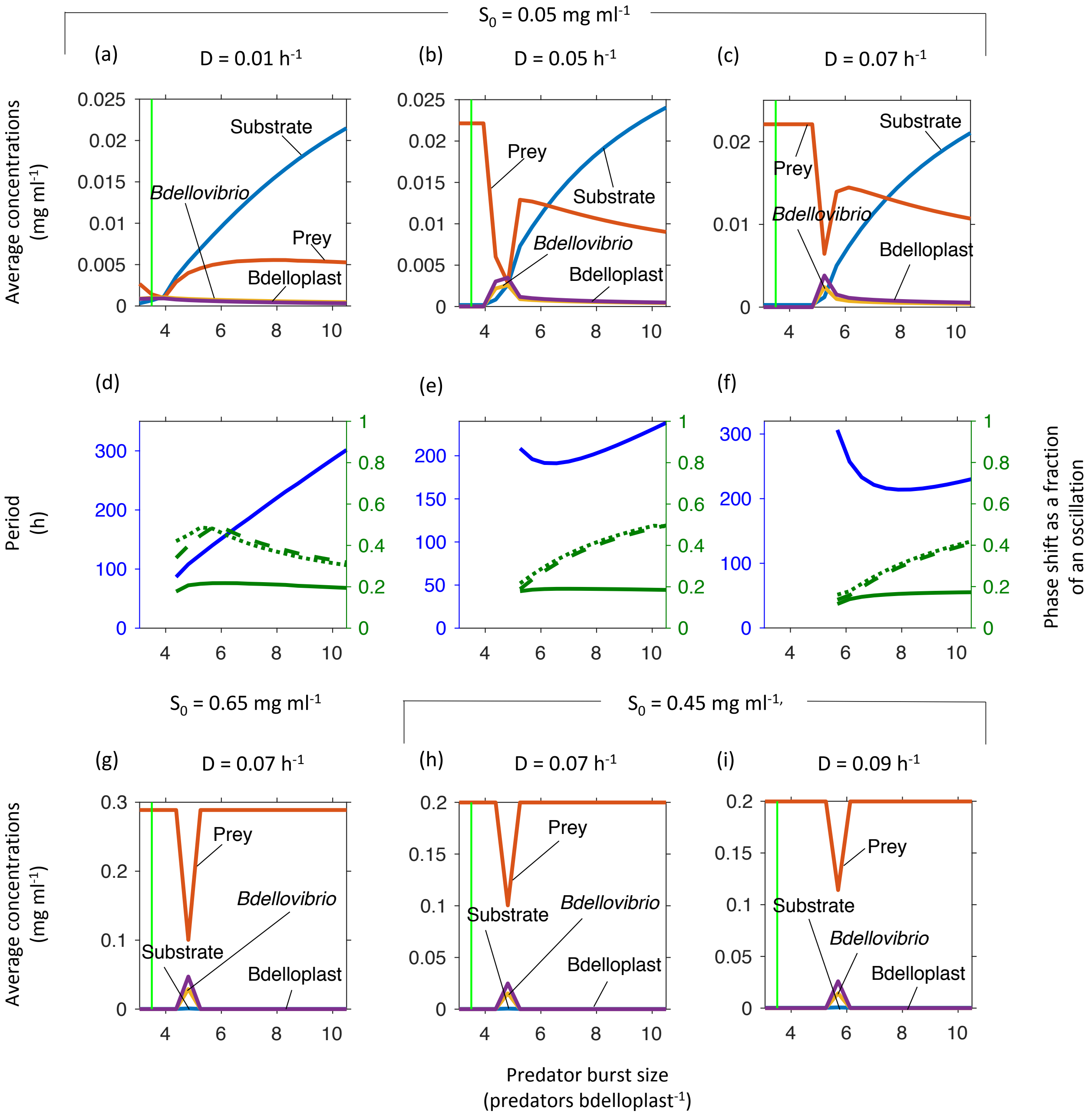

**Fig. S11** Minimal and optimal burst sizes. Oscillations occur above the optimal burst size, but only at low inflow substrate concentrations ($S_0$) (top row) not higher $S_0$ (bottom row). Top and bottom rows show concentrations at steady state or averaged over one oscillatory cycle. The burst size for an *E. coli* prey cell (3.5 predators bdelloplast$^{-1}$) is indicated by the vertical green line. Middle row shows the oscillatory period (blue, left axis) and phase shifts (green, right axis) from substrate peak to peak of prey (solid line), free *Bdellovibrio* (dashed line) or bdelloplast (dotted line) for the scenarios in the top row.

##### ***Why bacteriophage outcompetes Bdellovibrio***

Under both inflow substrate concentrations ($S_0$) tested, the bacteriophage outcompeted *Bdellovibrio* (Fig. 5). Compared to *Bdellovibrio*, the bacteriophage is hampered by an increased predator $K$-value ($K_{N,P}$), but helped by a lack of mortality, higher attack rate constant ($\mu_P$) and improved kinetics of prey consumption (increased $k_P$ and burst size). To find out which of these advantage(s) allowed the phage to outcompete *Bdellovibrio*, we ran competitions where the phage kept the $K_{N,P}$ disadvantage and had one or more of the advantages. We found that increased burst size ($Y_{P/B}$) alone was sufficient to allow the phage to outcompete the *Bdellovibrio* (Fig. S12d). Increased attack rate constant ($\mu_P$) and reduced mortality ($m$) were also sufficient, but only in combination (Fig. S12a, b, e). Increased maturation rate ($k_P$) was insufficient even in the presence of either an increased $\mu_P$ or reduced $m$ (Fig. S12c, f, g).

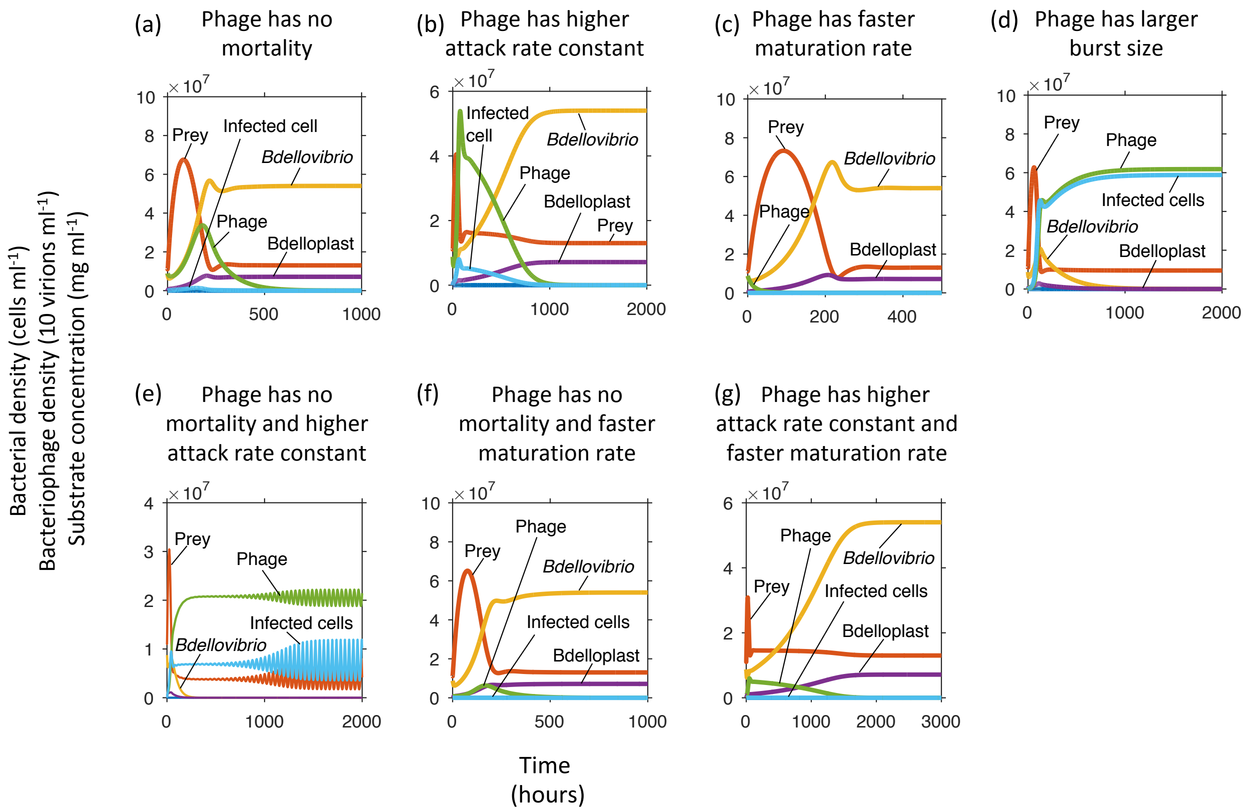

**Fig. S12** Competitions between *Bdellovibrio* and a phage that has one disadvantage, an increased predator $K$-value (higher $K_{N,P}$), but one or more compensating advantages. The phage wins if it has a larger burst size (panel **d**) or a combination of immortality and higher attack rate constant (panel **e**). All competitions were carried out at a dilution rate of 0.02 $h^{-1}$ and an inflow substrate concentration ($S_0$) of 0.05 mg $ml^{-1}$. To enable all data to be shown on the same axis, the phage data values were divided by 10, so 1 axis unit represents 10 virions.

***Predator spiking in a batch system***

Williams and co-workers spiked seawater mesocosms with *Vibrio parahaemolyticus* and observed that the naturally resident *Halobacteriovorax* (a marine predatory bacterium with the same lifecycle as *Bdellovibrio*) spiked in numbers, whilst no resident bacteriophages did (26). We used our model, with the dilution rate set to 0 $h^{-1}$ to mimic the batch culture setting of the mesocosm, to explore the response of *Bdellovibrio* and bacteriophage to a spike in prey numbers. We observed that in the absence of bacteriophage, *Bdellovibrio* densities spiked in response to the spike in prey (Fig. S13a). When both *Bdellovibrio* and bacteriophage where present, however, only the bacteriophage spiked in numbers – as expected given our results on *Bdellovibrio* versus phage competitions. Since this is in contrast to the findings in (26), we suspect that the mesocosms did not contain any phages able to infect the non-indigenous strain of *V. parahaemolyticus* added to the microcosms. The presence of such phages was not demonstrated.

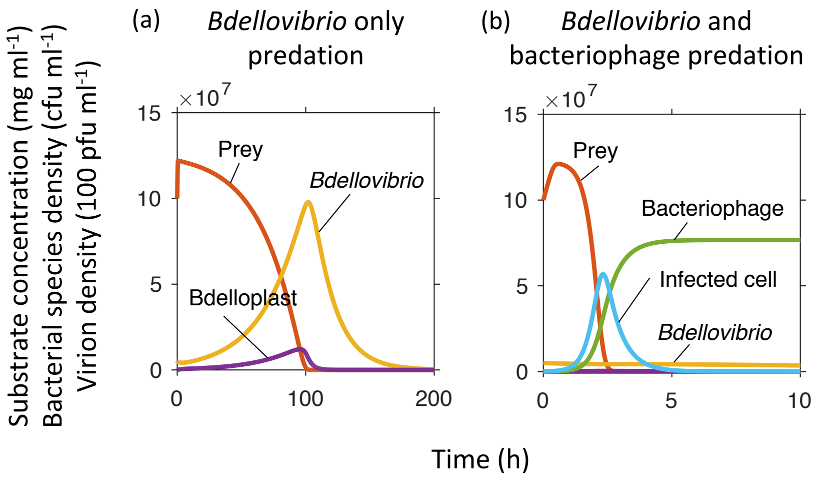

**Fig. S13** Predator response to a spike in prey in a closed batch system (dilution rate = 0 $h^{-1}$).

***Co-existence of two predators on one prey***

We found that a predator with a higher attack rate constant ($\mu_P$) and another with a lower predator $K$-value (lower $K_{N,P}$) could co-exist on the same prey assuming that there is a trade-off between these two parameters (Fig. S14). Depending on the extent of each advantage, the predators either co-existed in a steady state (Fig. S14a) or sustained oscillations (Fig. S14d). There were two combinations of $\mu_P$ and $K_{N,P}$ that allowed co-existence (Fig. S14e). Low inflow substrate concentrations ($S_0$) limited the maximal prey density, hence, the $K_{N,P}$ became the key factor (Fig. S14a, b). At higher $S_0$ there was more prey, so predators encountered prey more often and the $\mu_P$ became the key factor (Fig. S14c, d).

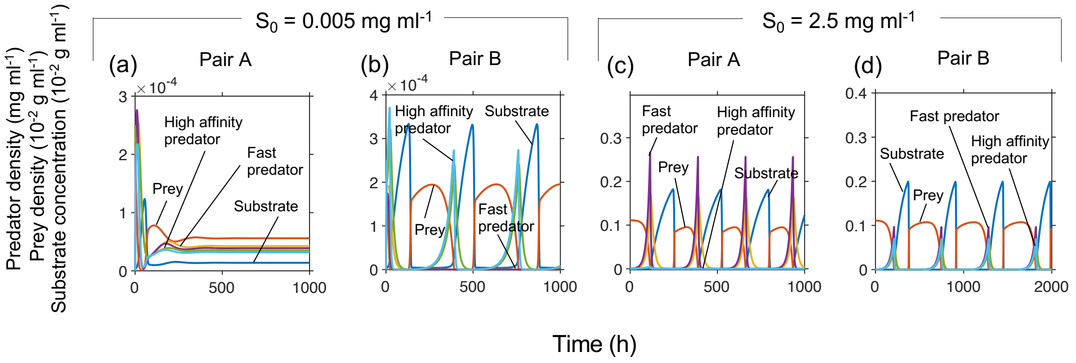

(e)

| | | $\mu_P$ (mg bdplast mg pred$^{-1}$ h$^{-1}$) | $K_{N,P}$ (mg dry mass prey ml$^{-1}$) |
|---|---|---|---|
| Pair A | Fast | 0.6 | 8.6 x 10$^{-4}$ |
| | Low half saturation constant | 0.38 | 3.4 x 10$^{-4}$ |
| Pair B | Fast | 0.39 | 8.6 x 10$^{-4}$ |
| | Low half saturation constant | 0.38 | 1.5 x 10$^{-5}$ |

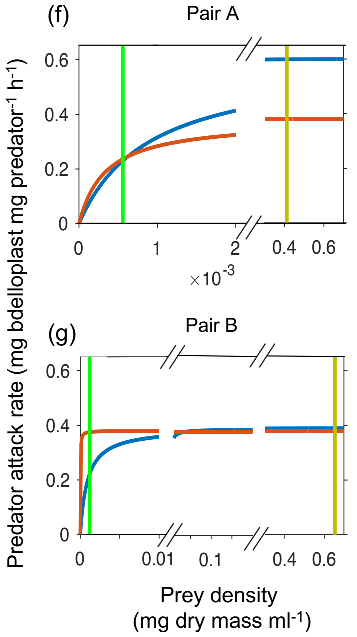

**Fig. S14** Due to a trade-off, a fast attacking predator (high $\mu_P$) can co-exist with a low $K$-value predator (low $K_{N,P}$). Dilution rate 0.02 h$^{-1}$. **a, c** Competition between predator pair A at two inflow substrate concentrations ($S_0$). **b, d** Competition between predator pair B at the same $S_0$ values. **e** Details of predator attack rate kinetics for both predator pairs. **f, g** Attack rates for predator pairs A (panel **f**) and B (panel **g**), fast predator (blue) and low $K$-value predator (red), over a range of prey densities. The green line indicates average prey density at low $S_0$ and the gold line at high $S_0$.

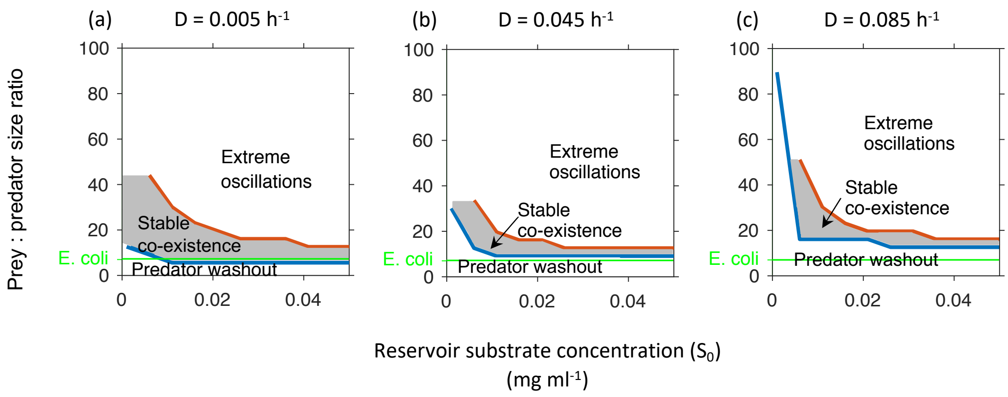

**Fig. S15** Paradox of enrichment. The range of prey to predator biomass ratios enabling permanence or stable co-existence (here including the regions of stable co-existence and damped oscillations in Fig. 2a) shrank with increasing productivity (inflow substrate concentration - $S_0$ ). At higher dilution rates, prey had to be unrealistically large for permanence; also, the prey biomass range for permanence shrank. Below the blue line, the predator is washed out. Above the red line, extreme oscillations occur resulting in bottlenecks and predator extinction due to stochastic dynamics. Within the shaded area, permanence (stable, long-term co-existence) occurred. The green line is the biomass ratio between an average *E. coli* and *Bdellovibrio* and similar to most prey used in laboratory studies or for isolating *Bdellovibrio* from the environment.

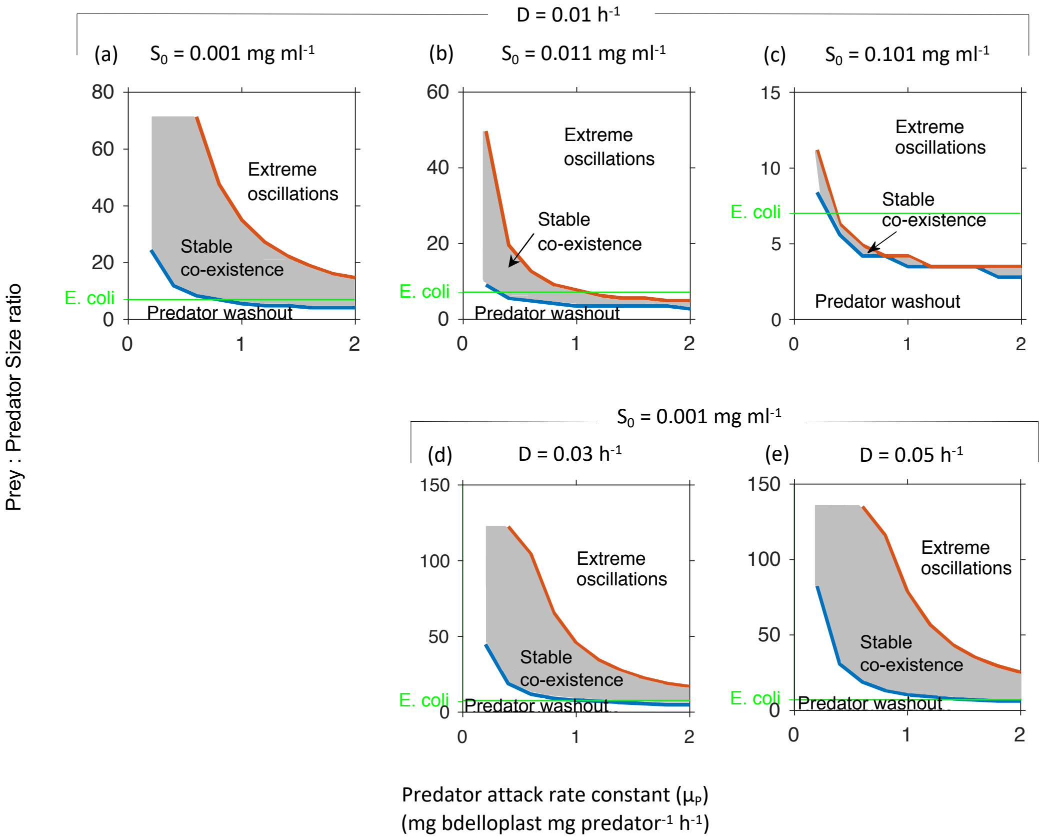

**Fig. S16** Tragedy of the commons. The prey biomass range for permanence of the predator (robust co-existence with prey) shrinks rapidly as the predator's attack rate constant ($\mu_P$) increases, as a too effective predator overexploits the prey, and then becomes extinct. This large drop in survival range occurs over a small increase in $\mu_P$, illustrating the sensitivity of the system to $\mu_P$ (see Fig. 3). Increased inflow substrate concentration ($S_0$) narrowed the prey biomass range for co-existence (top row), whilst increased dilution rate ($D$) expanded the range (panel **a** and bottom row). Below the blue line, the predator is washed out. Above the red line, extreme oscillations occur resulting in bottlenecks and predator extinction due to stochastic dynamics. Within the shaded area, permanence occurred. The green line is the biomass ratio between an average *E. coli* and *Bdellovibrio* and similar to most prey used in laboratory studies or for isolating *Bdellovibrio* from the environment.

***Global parameter sensitivity analysis***

When parameterising our model, we based our values on literature data from the best-studied predator strain (*B. bacteriovorus* strain HD100) predating the best-studied prey (*E. coli* MG1655) growing on glucose as the sole carbon and energy source (Table 1). Nevertheless, some processes such as prey growth are better studied, whilst others such as attack rate constant ($\mu_P$) are hardly studied. To understand how much the system behaviour would change if the parameters were different due to changes in substrate, prey or predator species, we conducted a global sensitivity analysis for each of the parameters identified from the dimensional analysis.

Note it was not possible to vary one of the dimensionless parameters ($Y'_{B*P}$) by changing a single dimensional parameter without affecting a second dimensionless parameter. Hence, we chose to vary each of its components ($Y_{B/_P}$ and $Y_{P/_B}$) in turn, resulting in a total of eight parameters being varied. Each parameter was increased and decreased by 1% for 10,000 settings of the other parameters over the ranges given in Table 1 and the relative change of substrate concentration or population densities was calculated.

For most of these parameter sets, a small change of the parameter value caused a correspondingly small change in substrate concentration or population densities or no change at all (Fig. S17). However, occasionally a 1% change in a parameter led to extreme changes in model output of several orders of magnitude (Fig. S18). These extreme sensitivities occurred where parameter sets were close to the boundaries of the dynamical regimes, e.g., between survival and extinction (cf. Fig. 2a).

Substrate concentration was only sensitive to the inflow substrate concentration (proportional response, Figs. S17a, S18a). Prey density was not sensitive to substrate concentration and prey growth kinetics (Monod parameters) but sensitive to predator parameters, particularly attack rate constant and biomass burst size where the response was more than proportional (Figs. S17b, S18b). Predator densities were also particularly sensitive to the attack rate constant but also to the maximal specific prey growth rate and less so to the yield of bdelloplast per predator, $Y_{B/_P}$ (Figs. S17c, S18c). Predator and bdelloplast densities responded similarly to parameter changes apart from the two parameters governing the conversion between bdelloplast and predator, i.e., bdelloplast maturation rate $k_P$ and yield of bdelloplast per predator, $Y_{B/_P}$ (Figs. S17c, d, S18c, d).

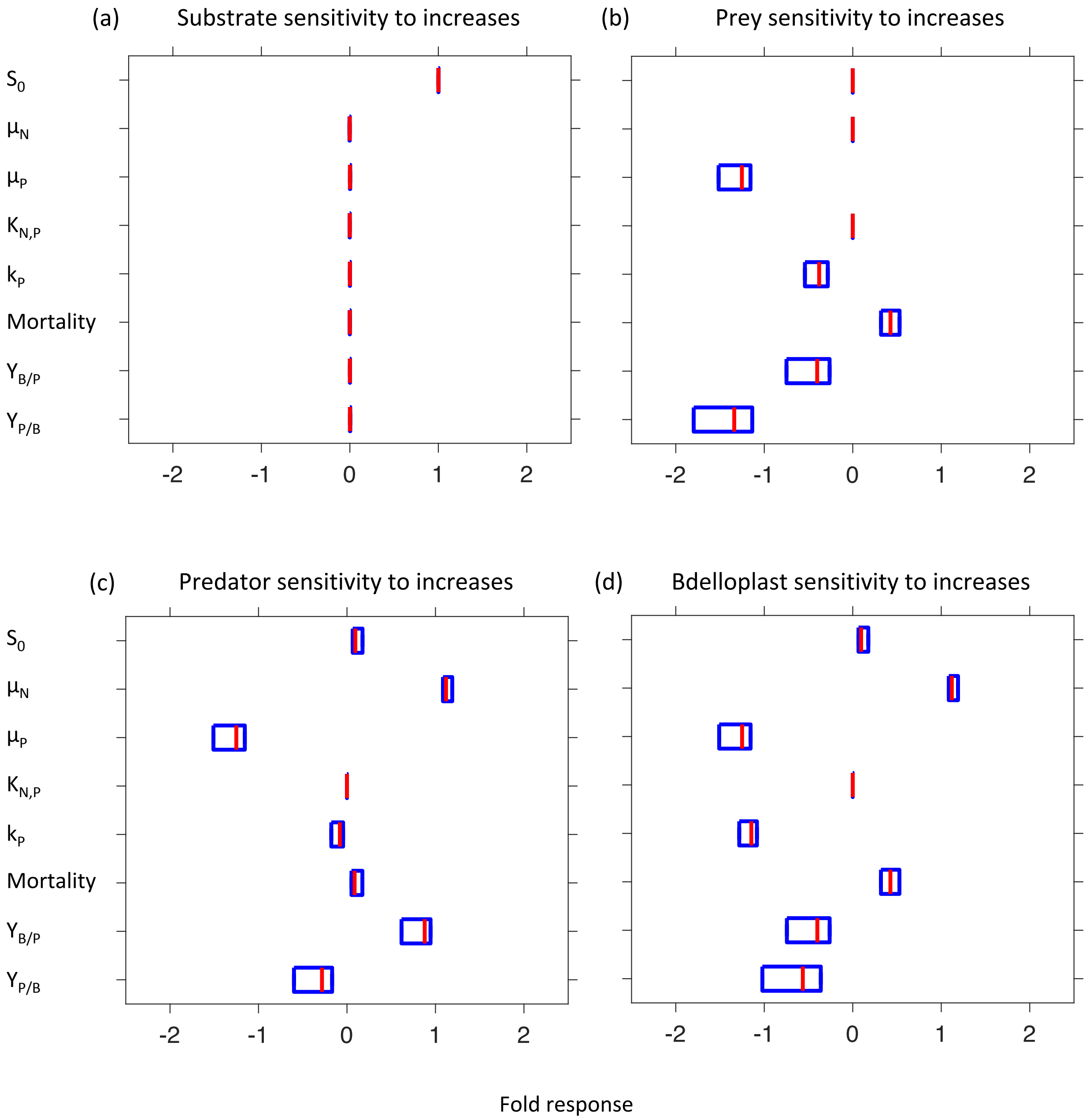

**Fig. S17** Global sensitivity of **a** substrate concentration, **b** prey, **c** predator and **d** bdelloplast population densities to small changes of a given model parameter (at 10,000 settings of the other model parameters). The distributions of the 10,000 sensitivities of outcomes to a 1% increase of the given model parameter are shown as box plots. Here, only median (red line) and 25th and 75th percentiles (blue lines) are shown; Fig. S18 includes the sometimes orders of magnitude different sensitivities outside the box.

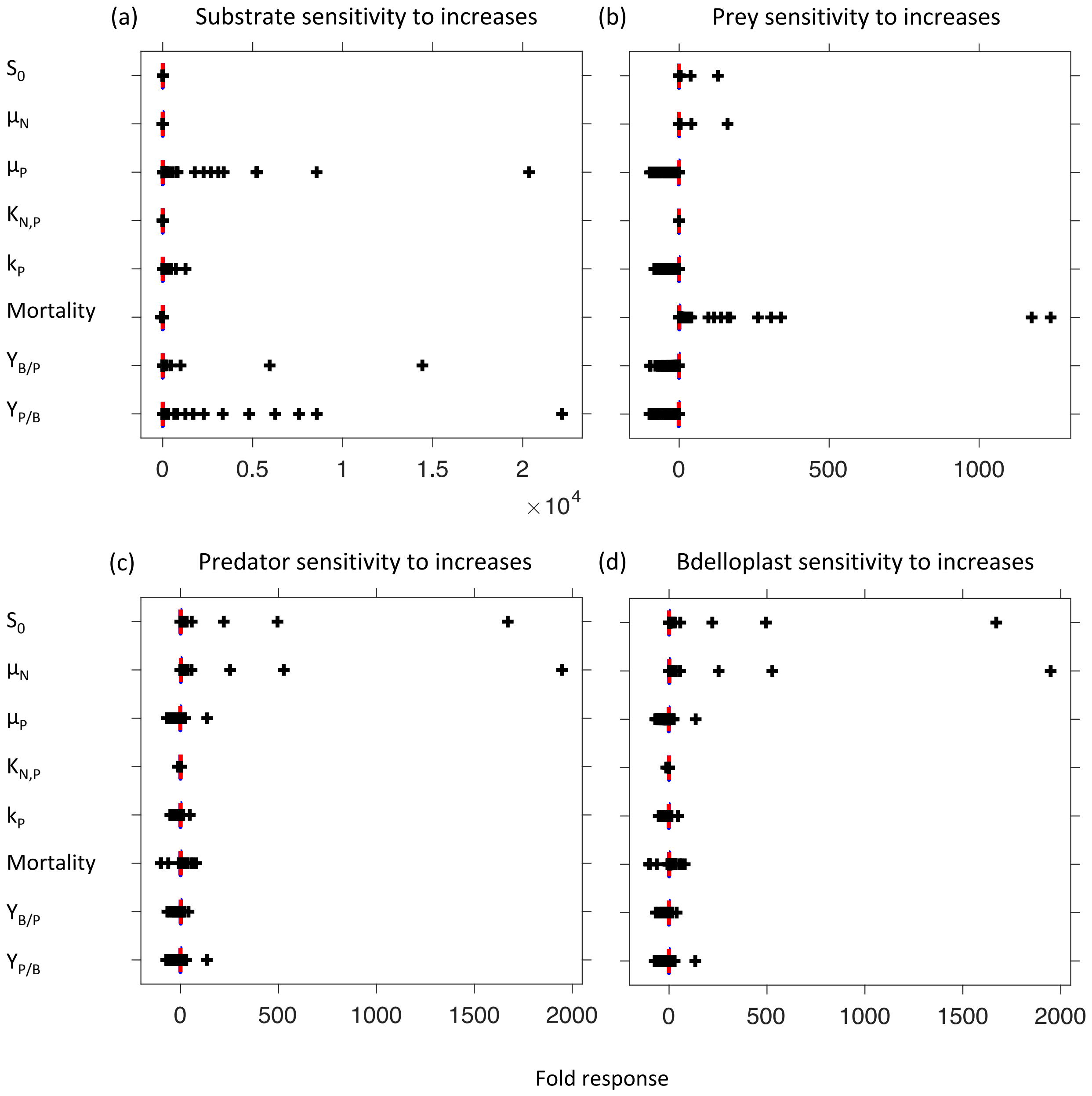

709 **Fig. S18** Global sensitivity analysis of model parameters as in Fig. S17 but now including all
710 values. At this scale, the blue 25th and 75th percentiles are indistinguishable from the median
711 in red. Black crosses are sensitivities outside the box and sometimes orders of magnitude
712 higher.

### ***References***

22.

19. Mckay MD, Beckman RJ, Conover WJ. 1979. Comparison of three methods for selecting values of input variables in the analysis of output from a computer code a comparison of three methods for selecting values of input variables in the analysis of output from a computer code. Technometrics 21:239–245.

20. Hadas H, Einav M, Fishov L, Zaritsky A. 1997. Bacteriophage T4 development depends on the physiology of its host *Escherichia coli*. Microbiology 143:179–185.

21. Stent GS, Wollman EL. 1952. On the two-step nature of bacteriophage adsorption. Biochim Biophys Acta 8:260–269.

22. Edelstein-Keshet L. 2005. Mathematical models in biologyClassics in Applied Mathematics. Society for Industrial and Applied Mathematics.

23. Rosenzweig ML, MacArthur RH. 1963. Graphical representation and stability conditions of predator-prey Interactions. Am Nat 97:209–223.

24. Curds CR, Bazin MJ. 1980. Protozoan predation in batch and continuous cultures, p. 115–176. *In* Droop, MR, Jannasch, HW (eds.), Advances in Aquatic Microbiology. Academic Press.

25. Rosenzweig ML. 1971. Paradox of enrichment: destabilization of exploitation ecosystems in ecological time. Science 171:385–7.

26. Williams HN, Lymperopoulou DS, Athar R, Chauhan A, Dickerson TL, Chen H, Laws EA, Berhane T-K, Flowers ARA, Bradley N, Young S, Blackwood D, Murray J, Mustapha O, Blackwell C, Tung Y, Noble RT, Williams HN, Florida A, King SML, Blvd J. 2016. *Halobacteriovorax*, an underestimated predator on bacteria: potential impact relative to viruses on bacterial mortality. ISME J 10:491–499.
